# Nuclear myosin VI regulates the spatial organization of mammalian transcription initiation

**DOI:** 10.1101/2020.04.21.053124

**Authors:** Yukti Hari-Gupta, Natalia Fili, Ália dos Santos, Alexander W. Cook, Rosemarie E. Gough, Hannah C. W. Reed, Lin Wang, Jesse Aaron, Tomas Venit, Eric Wait, Andreas Grosse-Berkenbusch, J. Christof M. Gebhardt, Piergiorgio Percipalle, Teng-Leong Chew, Marisa Martin-Fernandez, Christopher P. Toseland

## Abstract

During transcription, RNA Polymerase II (RNAPII) is spatially organised within the nucleus into clusters that correlate with transcription activity. While this is a hallmark of genome regulation in mammalian cells, the mechanisms concerning the assembly, organisation and stability which underpin the function these transcription factories remain unknown. Here, we have used combination of single molecule imaging and genomic approaches to explore the role of nuclear myosin VI in the nanoscale organisation of RNAPII. We reveal that myosin VI acts as the molecular anchor that holds RNAPII into transcription factories. Perturbation of myosin VI leads to the disruption of RNAPII localisation, changes in chromatin organisation and subsequently a decrease in gene expression. Overall, we uncover the fundamental role of myosin VI in the spatial regulation of gene expression during the rapid response to changes in the cellular environment.

## INTRODUCTION

The tight regulation of gene expression is critical for the maintenance of cellular homeostasis. This is fundamental during organism development and for the prevention of disease. In eukaryotic cells, RNA polymerase II (RNAPII) directs the flow of genetic information from DNA to messenger RNA (mRNA). Detailed genetic and biochemical assays have revealed a multi-level regulation of transcription, including *cis* control elements within the DNA and *trans* factors, such as general transcription factors, activators, repressors and a large number of coactivators. More recently, the actin-based molecular motors, myosins, have been also shown to act as transcription regulators (de Lanerolle, 2012, de Lanerolle and Serebryannyy, 2011, Fomproix and Percipalle, 2004, Hofmann et al., 2006, Kukalev et al., 2005). Myosins modulate their interaction with actin through their ATPase activity, which occurs within their highly conserved motor domain (Figure 1A). They are involved in multiple cellular processes including cell migration, endocytosis and exocytosis (Fili and Toseland, 2019). More recently, they have also been identified within the cell nucleus, where they have roles in transcription, DNA damage and chromosome organisation (de Lanerolle, 2012).

**Figure 1.**
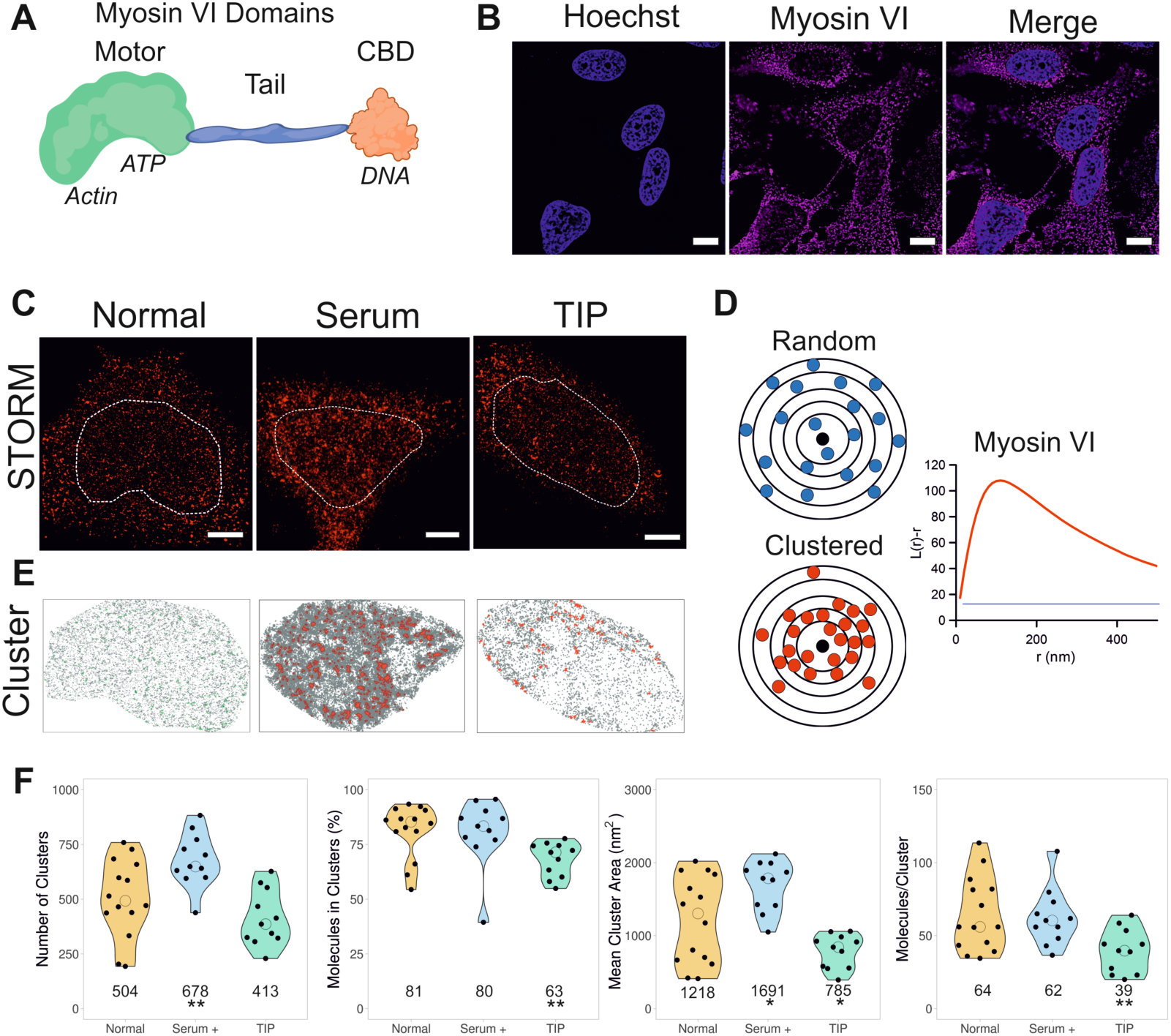
Nuclear organisation of myosin VI. (A) Cartoon depiction of the myosin VI domains and key features discussed in the text. The motor domain has ATPase activity regulates actin binding within the domain. The motor is connected to a cargo binding domain (CBD) in the myosin tail which has the ability to bind DNA. (B) Widefield Immunofluorescence staining against myosin VI (magenta) and DNA (cyan) in HeLa cells. Images were acquired at the mid-point of the nucleus (Scale bar 10 µm). (C) Example STORM render images of myosin VI under normal, serum- and TIP-treated conditions, as described in the Experimental Procedures (scale bar 2 µm). Dotted lines represent a region of interest (ROI) containing the nucleus which are taken forward for cluster analysis. The nucleus was identified using either Hoechst or RNAPII staining. (D) Depiction of molecular clustering and random distribution. We performed cluster analysis using the linearized form of Ripley’s K function (Pageon et al., 2016) *L(r)-r,* where r is the radius. A plot of *L(r)-r* versus *r* gives a value of zero for a random distribution (blue line), but deviates from zero, towards positive values, due to molecular clustering (red). The organisation of myosin VI is seen with a peak at 125 nm. (E) Cluster maps based upon the selected ROI in (c). Clusters are shown in green (Normal) or red (Serum and TIP treatment). (F) Cluster analysis of myosin VI nuclear organisation under normal, serum- and TIP-treated conditions. Individual data points correspond to the average value for a cell ROI (n = 14 for normal, 11 for serum- and TIP-treated). The values represent the mean from the ROIs for each condition (Only statistically significant changes are highlighted *p <0.05, **p <0.01 by two-tailed t-test compared to normal conditions).

The minus-end directed myosin, Myosin VI (Figure 1A), has been shown to bind DNA through its cargo binding domain (CBD) and couple itself to RNAPII in an actin-dependent manner through the motor domain (Fili et al., 2017). It has been revealed that the ability of myosin VI to bind DNA and its ATPase activity are both critical for transcription *in vitro* (Cook et al., 2018, Fili et al., 2017, Fili et al., 2020), and myosin VI can function in gene pairing (Zorca et al., 2015). Recently, myosin VI has been shown to actively undergo directed motion in the nucleus in response to transcription stimulation (Große-Berkenbusch et al., 2020). However, the precise role that this motor protein has in transcription has remained elusive.

The spatial organization of transcription has been debated and studied by both imaging and immunoprecipitation methods for over two decades (Cho et al., 2016a, Jackson et al., 1993, Papantonis and Cook, 2013). The formation of transcription centres has been suggested to increase the local concentration of enzymes and render these nuclear processes more efficient (Mao et al., 2011). Enzymatic clustering occurs in many cellular processes, particularly in the nucleus, with examples in replication (Kennedy et al., 2000) and DNA repair (Misteli and Soutoglou, 2009). Therefore, it is not surprising that clusters of RNAPII have been observed. The lifetime and composition of these clusters has been a matter of debate, with discrepancies between antibody staining in fixed cells, versus live cell observations (Jackson et al., 1993, Kimura et al., 2002, Papantonis and Cook, 2013, Sugaya et al., 2000, Sutherland and Bickmore, 2009, Zhao et al., 2014). More recently, RNAPII has been found to transiently cluster during transcription (Cho et al., 2016a, Cho et al., 2016b) and active RNAPII has been found to constrain chromatin dynamics (Nagashima et al., 2019). Yet, detailed molecular mechanisms of how these clusters form and how they are maintained remain unknown.

Interestingly, the biochemical properties of myosin VI can be tuned by the load applied to the motor (Altman et al., 2004), which allows it to switch from an active transporter to an actin anchor when tension is applied. We therefore hypothesised that myosin VI could act as either an anchor to stabilise RNAPII or as an auxiliary motor to drive RNAPII through the gene body. In either case, this would impact the organisation of RNAPII within the nucleus.

To this end, this study set out to explore whether myosin VI activity is responsible for the spatial organization of RNAPII. Through a combination of single molecule imaging and genomic studies, we have endeavoured to provide general mechanistic insight into how this form of nuclear organisation is achieved and what is the role of a myosin in this process.

## RESULTS

### The nuclear organisation of myosin VI

Myosin VI is present throughout the mammalian cell, including the nucleus (Figure 1B). To gain better understanding of the spatial organization of myosin VI, we used super resolution imaging, specifically Stochastic Optical Reconstruction Microscopy (STORM) (Figure 1C). Using this approach, individual myosin VI molecules within the nucleus could be resolved, quantified and their functional clustering behaviour assessed. To determine whether myosin VI assembles into clusters or is randomly distributed, we performed cluster analysis using the linearized form of Ripley’s K function (Pageon et al., 2016) (Figure 1D). This analysis demonstrated that nuclear myosin VI is clustered, rather than randomly distributed. To further understand this clustering behaviour, we used the Clus-DoC software (Pageon et al., 2016), which allows to quantify the spatial distribution of a protein by generating cluster maps (Figure 1E). We were able to determine that 81% (± 12) of nuclear myosin VI is clustered, with an average of 504 (± 178) clusters per nuclei. Each cluster, with an average cluster size of 1.2 µm^2^ (± 0.578), consists of 64 (± 24) myosin VI molecules, (Figure 1F). Of note, all the parameters quantified showed a large cell-to-cell variation, which may be attributed to the cells not being synchronized.

Given the well-established role of myosin VI in transcription (Cook et al., 2018, Fili et al., 2020, Fili et al., 2017, Große-Berkenbusch et al., 2020, Vreugde et al., 2006), we assessed whether stimulation of transcription can alter its clustering and thus functional properties. Indeed, stimulation with serum induced a significant increase in the nuclear distribution of myosin VI, as evidenced by both the STORM images and cluster maps (Figure 1C and E). Consistent with the noticeable increase in its nuclear recruitment, the number of clusters and their area increased significantly to 678 (± 122) clusters per nuclei and 1.7 µm^2^ (± 0.327), respectively (Figure 1F). However, the number of molecules per cluster and the overall percentage of molecules in a cluster remained unchanged, suggesting that new clusters are formed upon serum stimulation but that there may be an upper limit on the number of molecules within a cluster. Interestingly, the cell-to-cell variation decreased, potentially due to a more synchronised cellular response to serum.

Since myosin VI is an ATPase, we then explored whether its nuclear distribution and clustering behaviour was dependent upon its myosin motor activity. To this end, we used the small molecule inhibitor 2,4,6-triiodophenol (TIP) which is known to perturb the motor activity of myosin VI (Heissler et al., 2012) and impact upon transcription (Cook et al., 2018, Fili et al., 2020, Fili et al., 2017). TIP treatment disrupted the nuclear organisation of myosin VI (Figure 1C). STORM imaging and cluster analysis showed a significant decrease in all parameters, except for the number of clusters (Figure 1C, E-F). This suggests that inhibition of myosin VI motor activity interferes with its ability to assemble into clusters. This motor-dependent clustering behaviour indicates that actin could participate in the formation of these structures.

Having observed that transcription stimulation drives myosin VI cluster formation, we turned our attention to RNAPII to explore if there is a relationship between the clustering of both proteins. Firstly, we imaged RNAPII in the transcription initiation state (pSer5) using STORM and performed cluster analysis (Figure 2A). As it has previously been shown (Cho et al., 2016a, Jackson et al., 1993), we also observed clusters of RNAPII under normal growth conditions, whereby 42 (± 17) % of RNAPII was clustered, into 246 (± 140) clusters per cell, each containing 60 (± 27) molecules in an area of 2.7 (± 0.8) µm^2^, on average (Figure 2B). Interestingly, these clusters partially colocalized with the myosin VI clusters. In order to quantify this colocalization, we employed the Degree of Colocalisation (DoC) analysis which is available in the ClusDoC software (Pageon et al., 2016). The colour-coded co-localization cluster map highlights these regions, where, 15 (± 3) % of myosin VI and 22 (± 3) % of RNAPII colocalise (Figure 2C). The single-molecule nature of these measurements allowed us to further interrogate the data by comparing the features of colocalised and non-colocalised clusters. Whilst the number of colocalised clusters for both proteins is lower than the non-colocalised ones, these clusters are up to 2-fold larger in size for RNAPII and up to 10-fold larger for myosin VI, compared to the non-colocalised subpopulation (Supplementary Figure 1A and B). This also correlates with an approximate 50 % and 100 % increase in RNAPII and myosin VI molecules, respectively, within the colocalised clusters. We also determined that, in the colocalised clusters, there is a ratio of two myosin proteins for each RNAPII. Overall, this suggests there is synergy between the two proteins.

**Figure 2.**
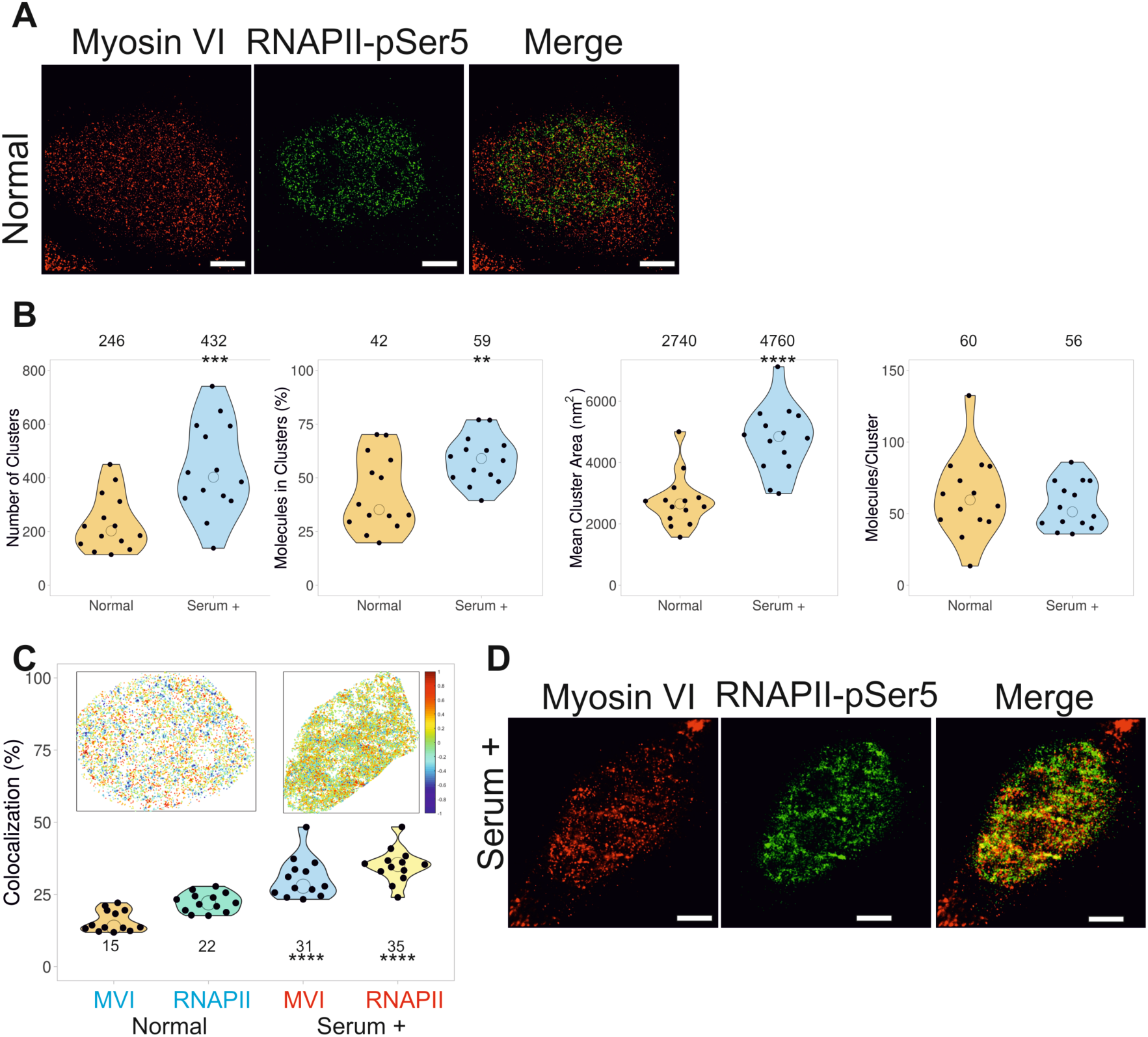
Nuclear organisation of RNAPII and colocalization with myosin VI. (A) Example STORM render image of myosin VI and RNAPII-pSer5 under normal conditions (scale bar 2 µm). (B) Cluster analysis of RNAPII nuclear organisation under normal and serum-treated conditions. Individual data points correspond to the average value for a cell ROI (n=14). The values represent the mean from the ROIs for each condition. (Only statistically significant changes are highlighted **p <0.01, ***p <0.001, ****p <0.0001 by two-tailed t-test compared to normal conditions). (C) Colocalisation analysis of myosin VI (MVI) and RNAPII-pSer5 clusters under normal and serum-treated conditions. Inset is a representative cluster colocalization heatmap whereby values of 1 are perfectly colocalised and −1 are separated from each other. Individual data points represent the percentage of each protein which is colocalized and correspond to the average value for a cell ROI (n=13). The values represent the mean from the ROIs for each protein (Only statistically significant changes are highlighted ****p <0.0001 by two-tailed t-test compared to normal conditions for each protein). (D) Example STORM render image of myosin VI and RNAPII-pSer5 under serum-treated conditions (scale bar 2 µm).

To further understand the functional relevance of the myosin VI-RNAPII co-localization, we stimulated transcription using serum (Figure 2D). Similar to myosin VI, there was a noticeable change in the RNAPII distribution. Serum stimulation led to an increase in cluster size and number of clusters of RNAPII, while the total number of molecules per cluster remained the same (Figure 2B). Moreover, we observed a significant increase in positive colocalization between the clusters, where 31 (± 7) % and 35 (± 6) % of myosin VI and RNAPII colocalized, respectively (Figure 2C). Furthermore, comparison of colocalised and non-colocalised subpopulations showed that the synergy between the two proteins is maintained following serum stimulation (Supplementary Figure 1A and B).

### Myosin VI regulates the spatial organisation of RNA Polymerase II

After observing the correlation between RNAPII and myosin VI clustering, and building upon the established role of myosin VI in transcription, we wanted to explore how myosin VI activity may impact the spatial organisation of RNAPII which underpins mammalian gene expression.

Treatment with the myosin VI inhibitor TIP induced a significant disruption of the spatial distribution and organisation of RNAPII, whereby the protein was aggregated at the nuclear periphery, while it was significantly decreased in the centre of the nucleus (Figure 3A and Supplementary Figure 2A). Not surprisingly, based on the visual redistribution of RNAPII, parameters such as the number, size, area and number of molecules per cluster were significantly decreased, compared to normal conditions (Figure 3B). We also confirmed that, while TIP impacts the clustering activity of myosin VI (Figure 1C), it does not lead to protein degradation (Figure 3C). We therefore conclude that the motor activity of myosin VI is required for RNAPII clustering. To further support this finding, we performed siRNA transient knockdown of myosin VI (Figure 3C, Supplementary Figure 2B and 3). Similar to the effect of the inhibitor, absence of myosin VI had a significant impact on the RNAPII distribution (Figure 3A). Based on the number of localisations in the STORM imaging and western-blot analysis, we confirmed that the total amount of RNAPII-pSer5 did not change following either treatment (Figure 3D and E). Therefore, perturbation of myosin VI causes destabilisation of RNAPII clusters. Consistently, TIP treatment also led to a decrease in myosin VI-RNAPII colocalization (Supplementary Figure 1C). Overall, our data indicated a role for myosin VI in the nuclear organisation of RNAPII.

**Figure 3.**
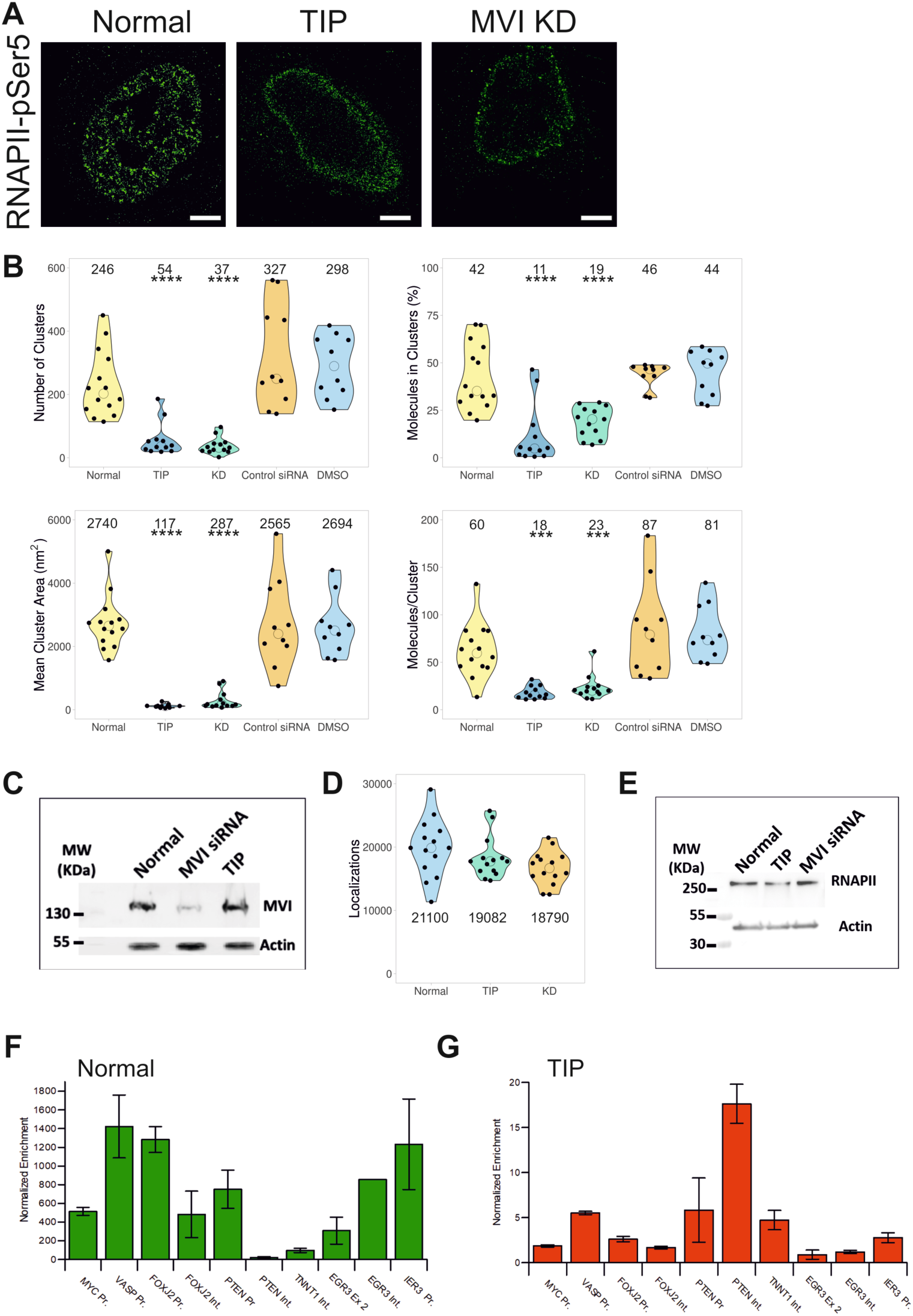
Spatial organisation of RNAPII depends upon MVI. (A) Example STORM render image of RNAPII-pSer5 under normal, TIP-treated and myosin VI (MVI) knockdown conditions (scale bar 2 µm). Further widefield example images are shown in Supplementary Figure 2A and B. (B) Cluster analysis of RNAPII nuclear organisation under the conditions described in (A). Individual data points correspond to the average value for a cell ROI (n=14 for normal, 12 for TIP, 13 for KD, 10 for Control siRNA and DMSO). The values represent the mean from the ROIs for each condition (Only statistically significant changes are highlighted ***p <0.001, ****p <0.0001 by two-tailed t-test compared to normal conditions). (C) Western-blot against MVI under normal, MVI-knockdown and TIP-treated conditions. Example widefield images of myosin VI under knockdown conditions are shown in Supplementary Figure 3. (D) Number of localisations of RNAPII-pSer5 under the different conditions. Individual data points correspond to the value for a cell ROI (n=14). The values represent the mean from the ROIs for each condition. (E) Western-blot against RNAPII-pSer5 under normal, TIP-treated and MVI knockdown conditions. (F) RNAPII-pSer5 ChIP against labelled loci. Values are the average of two independent experiments. Error bars represent SEM from two independent experiments. (G) RNAPII-pSer5 ChIP against labelled loci following TIP-treatment. Values are the average of two independent experiments. Error bars represent SEM from two independent experiments.

We next sought to establish whether the redistribution of RNAPII also correlated with its loss from chromatin. To this end, Chromatin Immunoprecipitation (ChIP) against RNAPII-pSer5 was performed under normal conditions and following TIP treatment. Indeed, myosin VI inhibition induced a decrease of several orders of magnitude in RNAPII occupancy from all tested loci (Figure 3F and G).

Both myosin VI knockdown and inhibition by TIP have an impact on the cytoplasmic and nuclear populations of myosin VI. To determine the specific role of the nuclear population of myosin VI in the spatial organisation of RNAPII, we transfected cells with NLS-tagged truncations of myosin VI, namely NLS-CBD (Cargo-binding domain) and NLS-Motor. Based on *in vitro* transcription assays (Fili et al., 2017), over-expression of these constructs and their targeting to the nucleus was expected to have a dominant negative impact upon the endogenous nuclear myosin VI by displacing the protein. Indeed, similar to TIP treatment and myosin knockdown, over-expression of either construct disrupted the nuclear distribution of RNAPII, as observed by widefield microscopy (Supplementary Figure 4A and B). We therefore concluded that it is the nuclear pool of myosin VI that is directly involved into the nuclear organisation of RNAPII.

**Figure 4.**
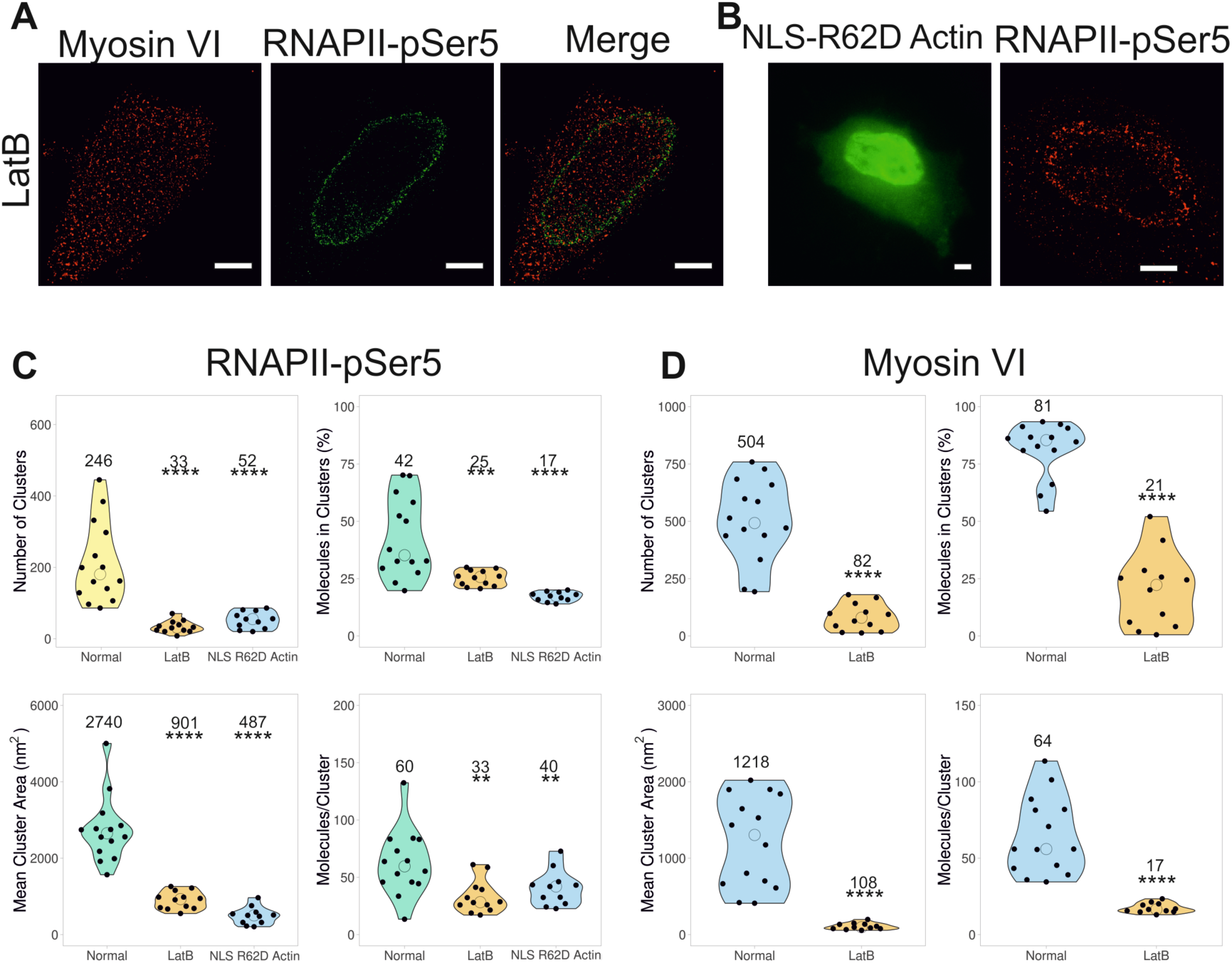
Impact of nuclear actin upon the organisation of RNAPII. (A) Example STORM render image of myosin VI and RNAPII-pSer5 following treatment with Latrunculin B (LatB), as described in the methods (scale bar 2 µm). Further widefield example images are shown in Supplementary Figure 2C. (B) (left) Example widefield image of YFP-NLS-R62D actin following transfection. (right) Example STORM render image of RNAPII-pSer5 following transfection of YFP-NLS-R62D actin (scale bar 2 µm). (C) Cluster analysis of RNAPII-pSer5 nuclear organisation following treatment with LatB (n=12) or transfection with YFP-NLS-R62D Actin (n=11). (D) Cluster analysis of myosin VI nuclear organisation following treatment with LatB (n=12). Individual data points correspond to the average value for a cell ROI. The values represent the mean from the ROIs for each condition (Only statistically significant changes are highlighted **p <0.01, ***p <0.001, ****p <0.0001 by two-tailed t-test compared to normal conditions).

We next explored whether the impact of myosin VI on RNAPII is also dependent upon nuclear actin. It has been well-established that actin is bound to RNAPII (de Lanerolle, 2012, Fomproix and Percipalle, 2004, Kukalev et al., 2005) and nuclear actin was recently found to support clustering of RNAPII (Wei et al., 2020). The association of myosin VI to RNAPII is also actin-dependent, *in vitro* (Fili et al., 2017). Moreover, as previously mentioned, the effect of TIP also suggests that actin filaments are involved. We therefore performed two types of actin perturbation experiments: (a) Treatment with latrunculin B to prevent actin polymerisation, which would reveal whether actin polymers are important for transcription. (b) Transient expression of a nuclear targeted monomeric actin mutant, namely NLS-YFP-R62D-actin (Serebryannyy et al., 2016). This mutant would bind to endogenous nuclear G-actin, thereby preventing its polymerisation. Both of these perturbations caused disruption of RNAPII organisation, revealing that polymerization of nuclear actin is critical to transcription (Figure 4A, 4B and Supplementary Figure 2C), as shown recently (Wei et al., 2020). Cluster analysis (Figure 4C) shows that all cluster parameters are significantly decreased in both conditions, as is the colocalization between myosin VI and RNAPII with latrunculin B (Supplementary Figure 1C). These results were also supported by the impact of latrunculin B treatment on the nuclear organisation of myosin VI. Similar to RNAPII, all clustering parameters for myosin VI were significantly decreased (Figure 4D), leading to a greater impact than TIP treatment (Figure 1). This suggests, that the nuclear roles of myosin VI involve its interaction with filaments or short polymers of actin. Overall, these results indicate that the myosin VI – actin interaction is required for the correct spatial organisation of RNAPII.

### Myosin VI controls the nuclear dynamics of RNA Polymerase II

Having revealed the role of myosin VI in the nuclear organisation of RNAPII, we then assessed its role in the dynamics of RNAPII in living cells to understand the assembly of the transcription factories.

To achieve this, we performed single molecule tracking of Halo-tagged or SNAP-tagged Rbp1, the largest RNAPII subunit, using an aberration-corrected multi-focal microscope (acMFM) system (Abrahamsson et al., 2013) (Figure 5A). This technique allows the simultaneous acquisition of 9 focal planes covering 4 µm in the *z* axis, with a 20 × 20 µm field of view, which is essentially the size of the HeLa cell nucleus. In this way, we were able to observe and track the 3D dynamics of RNAPII across the whole nucleus, in live cells (Figure 5B). Clustering of RNAPII was not observed in these live cell experiments due to the low labelling density required in order to achieve single-molecule detection in the crowded nuclear environment.

**Figure 5.**
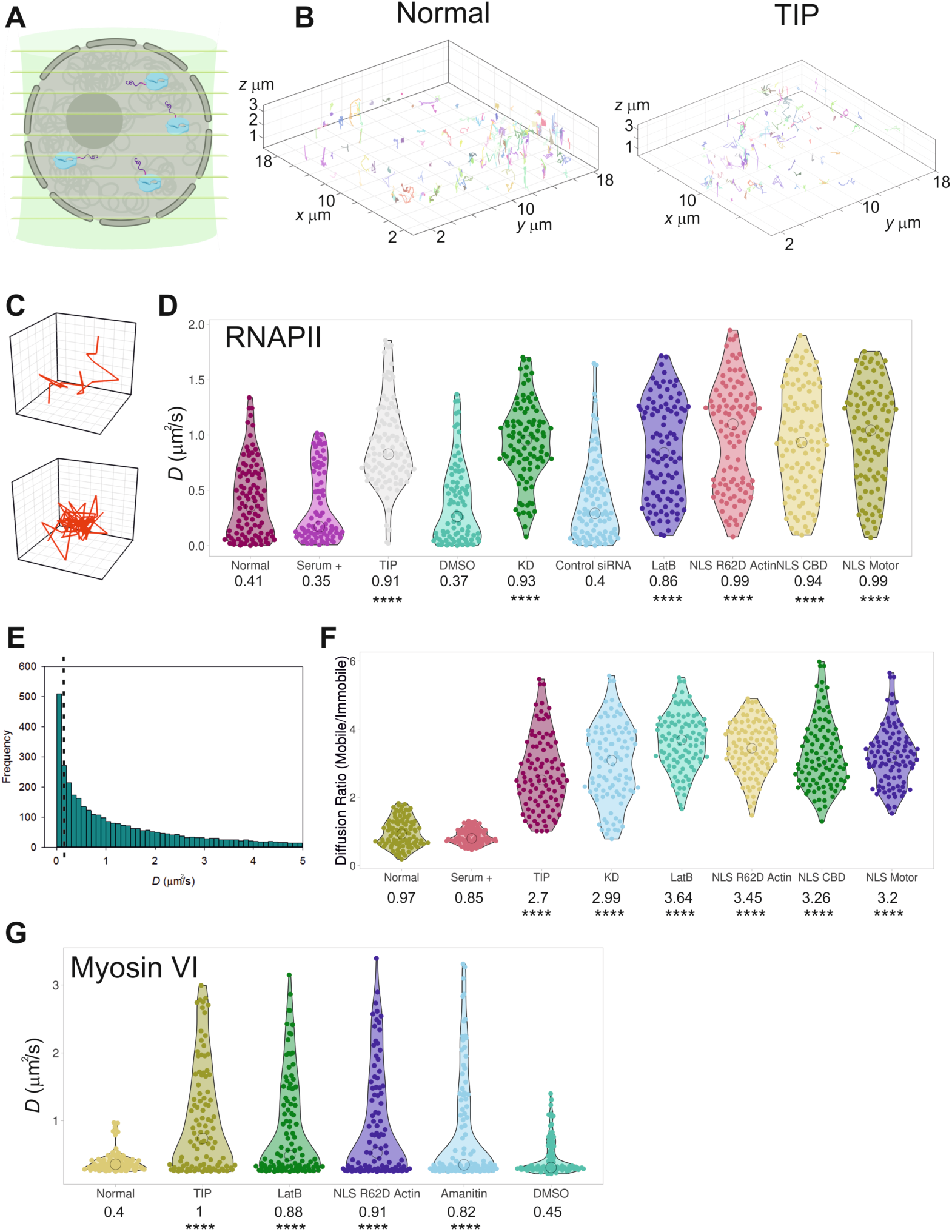
Live cell single molecule dynamics of RNAPII. (A) Cartoon depicting simultaneous acquisition of 9 focal planes covering 4 µm to perform live cell 3D single molecule tracking of RNAPII. (B) Example render of 3D single molecule trajectories under normal and TIP-treated conditions. (C) Example of trajectory of a diffusive and spatially confined molecule which can be identified in (B). (D) Plot of Halo-RNAPII or SNAP-RNAPII diffusion constants under the stated conditions derived from fitting trajectories to an anomalous diffusion model, as described in methods. NLS-R62D Actin, NLS CBD and NLS Motor refer to tracking of RNAPII following transfection of these constructs. NLS CBD and NLS motor were transfected in to SNAP-RNAPII cells. Individual data points correspond to the average value for a cell ROI (n = 100). The values represent the mean from the ROIs for each condition (Only statistically significant changes are highlighted ****p <0.0001 by two-tailed t-test compared to normal conditions). (E) Example histogram of diffusion constants arising from a single cell. The dotted line represents the threshold applied to segregate static and dynamic molecules. (F) Using the threshold defined in (e), trajectories were plotted as a ratio of mobile and immobile species. Individual data points correspond to the average value for a cell ROI. The values represent the mean from the ROIs for each condition (Only statistically significant changes are highlighted ****p <0.0001 by two-tailed t-test compared to normal conditions). (G) Plot of Halo-myosin VI diffusion constants under the stated conditions derived from fitting trajectories to an anomalous diffusion model, as described in methods. Individual data points correspond to the average value for a cell ROI (n = 100). The values represent the mean from the ROIs for each condition (Only statistically significant changes are highlighted ****p <0.0001 by two-tailed t-test compared to normal conditions).

Moreover, all populations of RNAPII were visible, compared to solely the pSer5 population in the STORM measurements. We observed pools of spatially confined RNAPII molecules and pools of molecules diffusing freely within the nucleus (Figure 5C). We determined the diffusion constant for each track by measuring the Mean Squared Displacement (MSD) and then plotted the average diffusion constant per cell (Figure 5D). Under normal conditions, we found that, on average, RNAPII diffuses relatively slowly (0.41 µm^2^ s^−1^) and this decreases further during transcription stimulation (0.35 µm^2^ s^−1^).

When we interrogate the individual tracks for each cell, we observed several populations of RNAPII which we termed as (i) static D<0.1 µm^2^ s^−1^, (ii) diffusive 0.1> D <5 µm^2^ s^−1^ and (iii) hyper mobile (not quantified) (Figure 5E). These findings are consistent with the previous reports using this technique (Abrahamsson et al., 2013). To further investigate RNAPII dynamics, the total numbers of static (<0.1 µm^2^ s^−1^) and mobile (>0.1 µm^2^ s^−1^) RNAPII molecules were plotted as a ratio (Figure 5F). Under normal conditions, just over half of the RNAPII population (52%) was static, probably corresponding to molecules confined at sites of transcription activation. Interestingly, this value was similar to the percentage of RNAPII molecules in clusters detected by STORM. Similar to the STORM experiments, we then observed the dynamics of RNAPII following TIP treatment, siRNA knockdown of myosin VI and actin perturbations. These treatments led to a 2-fold increase in the RNAPII average diffusion constant (Figure 5D) and 3 to 4-fold increase in the motile fraction (Figure 5F). Visually, the impact was also clear and could be observed through the loss of spatially confined molecules and the gain of diffusive tracks (Figure 5B). This was further quantified by plotting the anomalous diffusion alpha values whereby TIP treatment leads to an increase in freely diffusing species (Supplementary Figure 5A). In order to assess whether TIP has a global impact on molecular diffusion in the nucleus, we transiently expressed an isolated SNAP-tag domain to act as a diffusion reporter for the nuclear environment (Supplementary Figure 5B). We would not expect any impact on the diffusion of this isolated protein domain when cells are treated with TIP. Indeed, no changes were observed, confirming that the detected changes in RNAPII behaviour relate solely to the activity of myosin VI. We also observed the RNAPII dynamics in cells transiently expressing the dominant negative NLS-motor and NLS-CBD constructs. Consistent with the effect of TIP and myosin VI knockdown, there was an almost 2-fold increase in RNAPII diffusion in both cases, as well as an increase in the motile fraction of RNAPII (Figure 5D and F). This increased mobility of RNAPII following perturbation of myosin VI would be expected to lead to a decrease in the number of clusters, which is what we observed with the STORM measurements. Moreover, the greater mobility of RNAPII would also account for its relocation to the nuclear periphery, where it may non-specifically associate with the nuclear membrane or lamina. Overall, our observations suggest a model whereby myosin VI stabilises the RNAPII at sites of transcription initiation.

Finally, we also explored the nuclear dynamics of myosin VI and its interplay with RNAPII (Figure 5G). Overall, myosin VI is relatively static, with a mean diffusion constant of 0.4 µm^2^ s^−1^. However, treatment with TIP, or perturbation of actin, resulted in an increased mean diffusion to approximately 1 µm^2^ s^−1^, which is consistent with the STORM measurements, that show a reduction in clustering behaviour. Interestingly, a 2-fold increase in mean myosin VI diffusion was observed when cells were treated with the RNAPII inhibitor α-amanitin, that inhibits transcription through RNAPII degradation. The impact of RNAPII on myosin VI dynamics indicates a two-way communication between the two proteins, as it would be expected for two interacting molecules.

### RNA Polymerase II spatial distribution is coupled to transcription activity

We then explored the impact of the perturbed nuclear organisation of RNAPII on the underlying chromatin which could fundamentally alter the cellular properties. We performed a high-content screening assay, using antibodies against histones H3K9ac and H3K27ac, positive epigenetic marks of active gene expression, and H3K9me3, a mark of repressed transcription (Wang et al., 2008). Fluorescent intensity was used as a readout for the level of each marker in cells grown under normal conditions and upon treatment with TIP (Figure 6A). Treatment with TIP led to a decrease in active transcription markers by 35% and 10% for H3K9ac and H3K27ac, respectively, and an increase in the repressive marker H3K9me3 by 100% (Figure 6B). To assess the overall impact on cell function, we performed live-cell growth assays under normal and myosin VI knockdown conditions. A 3-fold decrease in growth was observed following knockdown (Figure 6C). The increase in growth rate after 3 days is consistent with the end of the transient knockdown. Overall, this change is indicative of a larger cellular response to the perturbation of RNAPII and the resulting decrease in gene expression.

**Figure 6.**
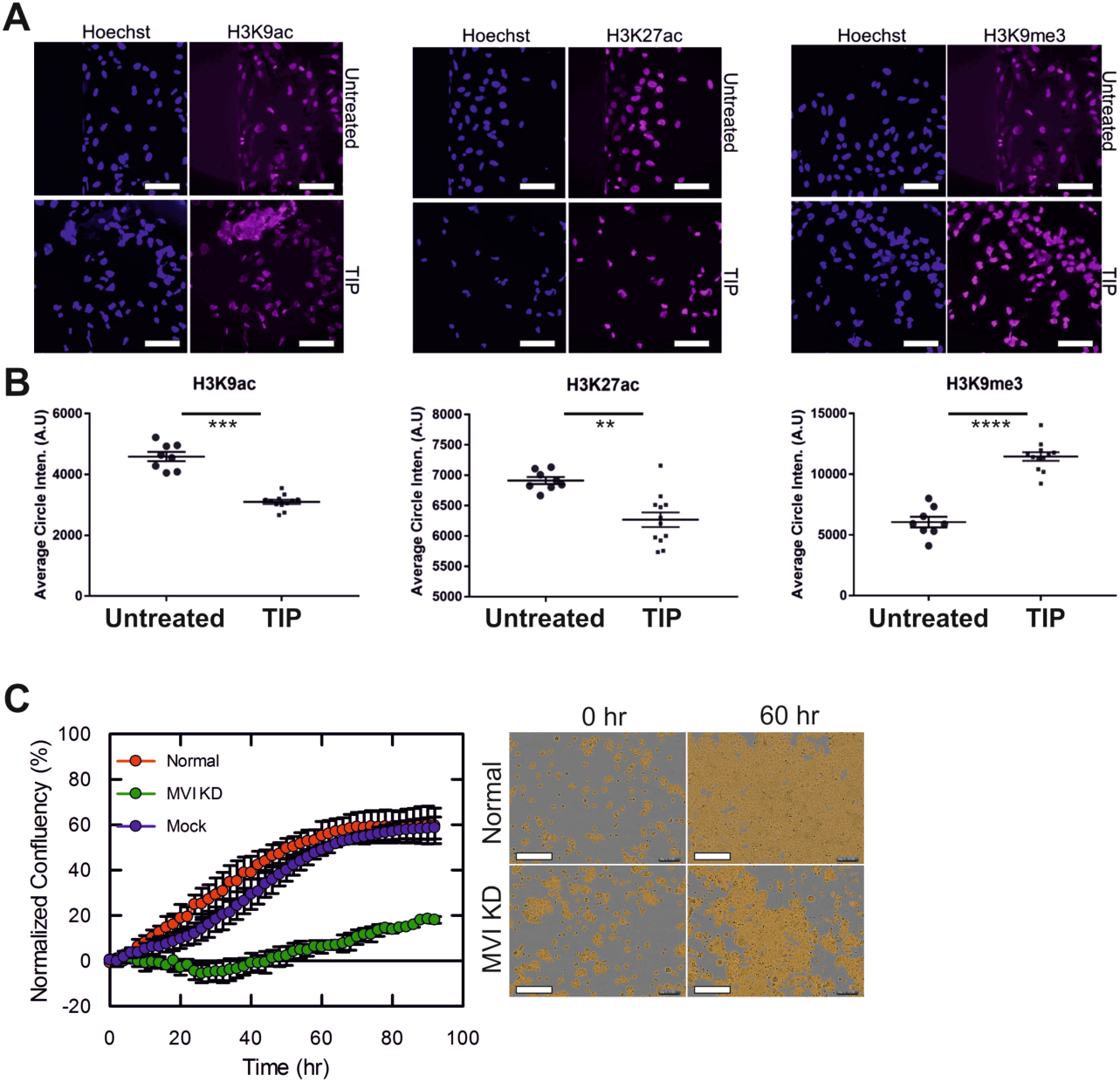
Perturbation of myosin VI impacts chromatin organisation and cell growth. (A) Example widefield Immunofluorescence staining against stated histones (magenta) and DNA (blue) in HeLa cells under normal and TIP-treated conditions (scale bar 100 µm). (B) The fluorescence intensity within the nucleus was measured for each histone marker under untreated and TIP-treated conditions, and each data point represents the average from a minimum of 1000 nuclei (Only statistically significant changes are highlighted **p <0.01, ***p <0.001, ****p <0.0001 by two-tailed t-test compared to untreated conditions). (C) Real-time growth of HeLa cells (red) and corresponding measurements following myosin VI (MVI) siRNA knockdown (green) and mock transfection control (blue). Data represent three independent measurements and error bars show SEM. Example images at start and 60 hr time points are shown (scale bar 300 µm in all images).

To explore the global changes in gene expression, RNA-seq measurements were performed under normal and myosin VI knockdown conditions. In total, we observed a significant change in the expression of 1947 genes (Log_2_ FC >0.5 or <0.5 with adjusted p value <0.05). From this set, 489 genes were up-regulated and 1458 were down-regulated (Figure 7A), which highlights the extensive negative impact on transcription due to the disruption of myosin VI. This is consistent with the STORM data that demonstrated disruption of the spatial organisation of RNAPII under these conditions (Figure 3A and 3B). The down-regulated genes were taken forward to Gene Ontology analysis.

**Figure 7.**
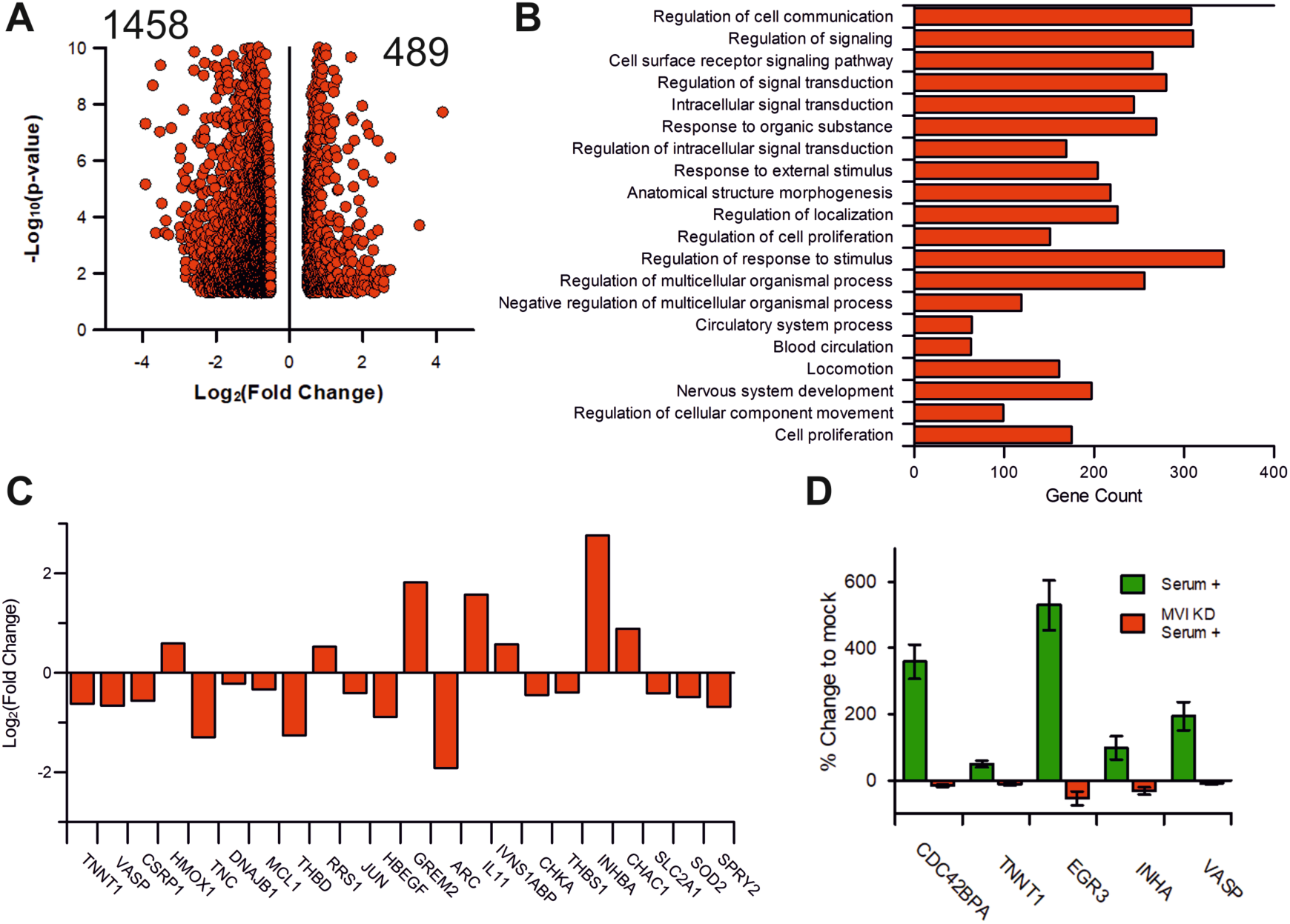
Impact of myosin VI perturbation on gene expression. (A) Volcano plot of differentially expressed genes following myosin VI knockdown. (B) GO terms, with gene count, for Biological Process corresponding to the genes negatively expressed following myosin VI knockdown. GO Terms are plotted based on significant enrichment, as shown in Supplementary Table 1. (C) Plot of gene expression changes for serum-responsive genes within the list of differentially expressed genes following myosin VI knockdown. (D) RT-qPCR Gene expression analysis of 5 serum-responsive genes with treatment of serum alone, or serum following a myosin VI (MVI KD) knockdown. Data are plotted relative to non-stimulated expression from three independent experiments. Error bars represent SEM from three independent experiments.

The breakdown for GO Biological Process reveals that affected genes are significantly enriched to processes such as “Regulation of signalling”, “Regulation of cell communication”, “Response to stimulus” and “Regulation of cell proliferation” (Figure 7B and Supplementary Table 1). Overall, the majority of the processes affected are coupled to cell response pathways, rather than to housekeeping ones. Hence, disruption of myosin VI perturbs expression of specific genes, but it does not completely halt transcription.

We observed that transcription stimulation with serum had a significant impact on the nuclear organisation of both myosin VI and RNAPII (Figure 1 and 2). Interestingly, the RNA-seq data also revealed that, out of 22 serum-responsive genes, two thirds were down-regulated when myosin VI was perturbed (Figure 7C). To investigate the role of myosin VI under conditions of transcription stimulation, we performed serum stimulation on cells where myosin VI had been knocked down. We then used RT-qPCR to monitor the expression of serum responsive genes. Control measurements showed that serum stimulation increases the expression of CDC42BPA, TNNT1, EGR3, INHA and VASP genes (Figure 7D). In all cases, knockdown of myosin VI completely abrogated this response. Taken together, the data shows that perturbation of myosin VI, which impacts the spatial organisation of RNAPII, impedes gene expression under stimulatory conditions. Therefore, myosin VI is critical for the cell’s response to stimulus.

### Myosin VI acts as a molecular anchor

We have revealed that myosin VI is a key regulator of RNAPII spatial organisation. However, how mechanism governing how myosin VI achieves this fundamental role is unknown. Based on previously conducted biochemical analysis, we hypothesised that myosin VI is bound to chromatin and/or transcription regulators through its CBD, and to RNAPII through actin (Fili et al., 2017).

Myosin VI is a rare motor protein with the ability to switch from a motile state to a molecular anchor, when forces greater than 2pN are applied to the molecule (Altman et al., 2004). This property prevents the unwanted dissociation from actin by increasing the affinity of the interaction. RNAPII is a large macromolecular machine which could diffuse or potentially move along DNA, away from transcription initiation sites. Such a movement would apply load upon myosin VI and then trigger the motor protein to anchor RNAPII *in situ*. To test this hypothesis, we set out to disrupt the ability of myosin VI to respond to force. To achieve this, we inserted a molecular spring consisting of a repeated penta-peptide sequence from spider-silk flagelliform into the myosin VI tail (Figure 8A). The repeat sequence has been widely used as a calibrated tension sensor (Grashoff et al., 2010). The spring unfolds as tension up to 10 pN is exerted across the molecule (Grashoff et al., 2010), thereby preventing load-induced changes on myosin VI. Therefore, this myosin VI construct should not be responsive to force up to 10 pN.

**Figure 8.**
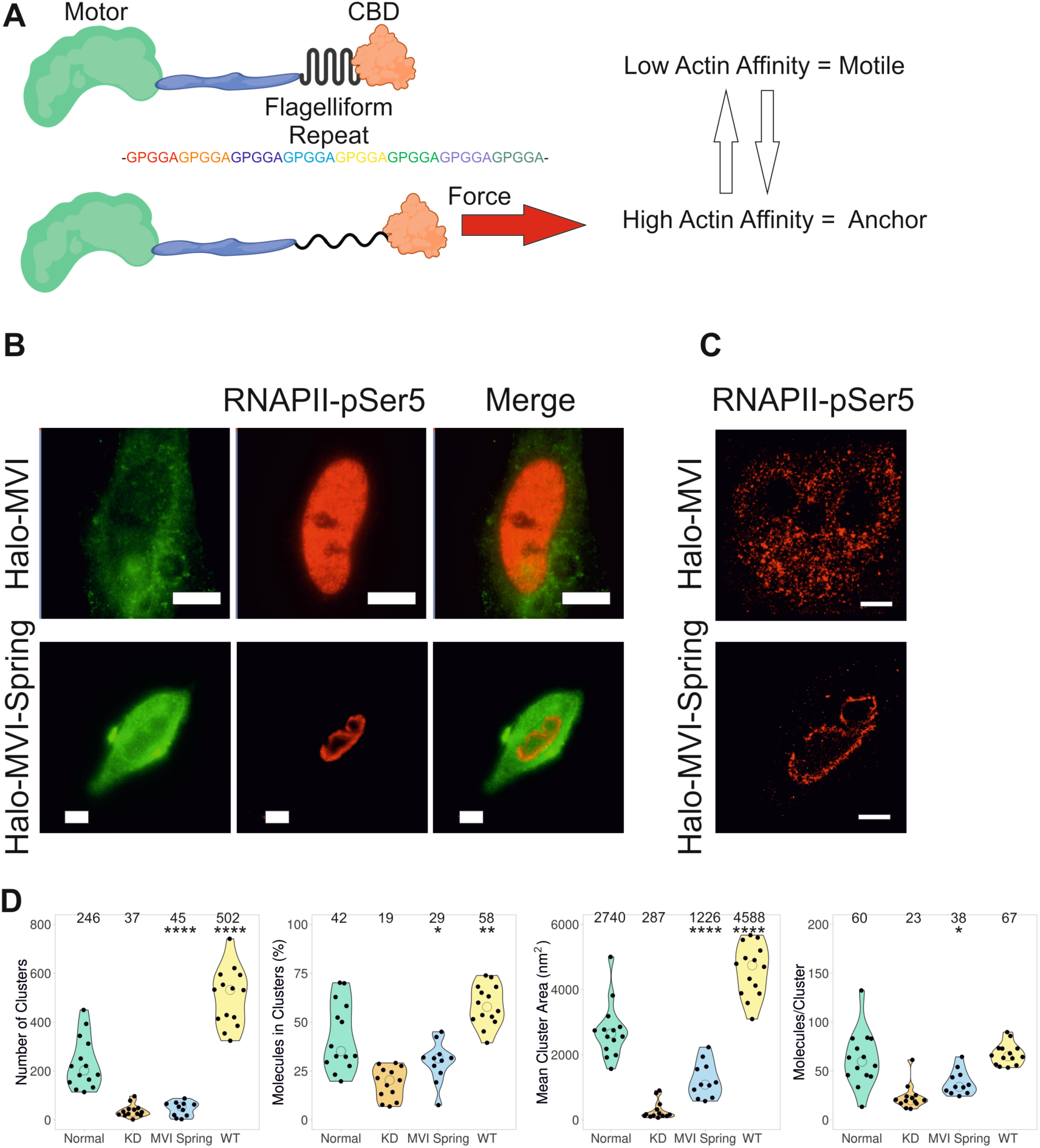
Myosin VI anchors RNAPII at transcription sites. (A) Cartoon depiction of myosin VI (MVI) containing the molecular spring (Flagelliform repeat) inserted proximal to the CBD. At low force, the spring is folded and myosin VI is in a low actin affinity mode. The application of force leads to extension of the spring which triggers the high affinity actin binding mode. (B) Example widefield imaging of Halo-MVI and Halo-MVI-spring stained with JF549 (green) and corresponding Immunofluorescence staining against RNAPII-pSer5 (red) in HeLa cells (Scale bar 10 µm). Further example images are in Supplementary Figure 6E. (C) Example STORM render image of RNAPII-pSer5 following transfection of Halo-MVI and Halo-MVI-spring, as described in the methods (scale bar 2 µm). (D) Cluster analysis of RNAPII-pSer5 nuclear organisation following treatment in (c). WT refers to Halo-MVI transfection. Individual data points correspond to the average value for a cell ROI. (n = 14 for Normal, 13 for KD, 11 for MVI Spring and 14 for WT) The values represent the mean from the ROIs for each condition (Only statistically significant changes are highlighted *p <0.05, **p <0.01, ****p <0.0001 by two-tailed t-test compared to normal conditions).

We first explored the impact of the insertion upon the biochemical properties of myosin VI. Firstly, CD spectroscopy confirmed that the recombinant protein is folded and stable (Supplementary Figure 6A), similar to wild type (WT) myosin VI (Supplementary Figure 6B). Moreover, the actin-activated ATPase activity was not affected by the presence of the insert, with *k*_cat_ 5.9 s^−1^ and 5.5 s^−1^ for myosin VI spring and WT, respectively (Supplementary Figure 6C). To assess whether the spring insert disrupts the load-induced anchoring ability in myosin VI, we then compared the ATPase rate of two stable dimeric constructs of the protein, one containing and one lacking the insert.

The stable dimeric myosin VI, in which a leucine zipper replaces the myosin VI C-terminal domain at residue 920 to dimerize the protein, has been widely used in myosin VI studies (Große-Berkenbusch et al., 2020, Mukherjea et al., 2014, Mukherjea et al., 2009, Park et al., 2006, Phichith et al., 2009). We first confirmed that the spring dimeric construct is indeed dimeric, as shown by a similar size-exclusion chromatography elution profile to wild type dimeric myosin VI (Supplementary Figure 6D).

As previously observed (Sweeney et al., 2007), the dimeric form of myosin VI displayed gating ATPase activity, whereby only one motor domain within the dimer hydrolyses ATP at a given point. Therefore, the observed ATPase rate was half of that of the monomeric myosin VI (*k*_cat_ 2.84 s^−1^), as seen in Supplementary Figure 6C. This gating behaviour results from load-induced conformation changes: the conformation of the leading motor is in a state which prevents ADP dissociation and therefore remains bound to actin (Altman et al., 2004). However, the insertion of the spring within the dimer construct showed an ATPase rate 4.9 s^−1^, which is similar to that of the monomeric myosin VI (Supplementary Figure 6C). This suggests the two motor domains are functioning independently. We propose that the spring-induced flexibility and the resulting inability to respond to load prevents the communication between the leading and the rear motor domains within the dimer.

Having assessed the biochemical properties of the spring construct, we transiently overexpressed it into mammalian cells where endogenous myosin VI had been knocked down (Figure 8B and Supplementary Figure 6E). The spring construct was localised throughout the cytoplasm and the nucleus. We then assessed the ability of the spring construct to rescue the disrupted RNAPII nuclear organisation, in comparison to the transiently overexpressed wild type myosin VI. Unlike wild type myosin VI (Figure 8B), the spring construct was unable to fully rescue the RNAPII nuclear distribution. As demonstrated by the STORM imaging and cluster analysis (Figure 8C and D), this partial rescue was evidenced in terms of number of clusters, cluster area and molecules per cluster. These results suggest that the ability of myosin VI to respond to force is required for rescuing the disrupted RNAPII distribution. In further support of this, the over-expression of wild type myosin VI was not only able to rescue RNAPII distribution, but to also increase RNAPII clusters number, size and percentage of molecules in clusters (Figure 8C and D). A possible explanation for the partial rescue by the spring construct could be that, at selected locations within the nucleus, the forces exerted upon myosin VI exceed the 10 pN limit, above which the spring insert would be fully unfolded and therefore responsive to forces and able to anchor. Therefore, we propose that the force-induced anchoring ability of myosin VI is critical for the nuclear organisation of RNAPII and that myosin VI physically holds RNAPII at sites of transcription initiation.

## DISCUSSION

Following a multidisciplinary approach, we have been able to shed light on to the regulation of transcription by addressing how the spatial organisation of transcription factories is achieved and the role of nuclear myosins in this process. We have observed that the molecular motor myosin VI is clustered within the nucleus and we showed that this activity is linked to the spatial organisation of RNAPII into transcription factories. We have also been able to show that the spatial and dynamic changes in RNAPII behaviour are dependent upon myosin VI and relate to wider chromatin and transcriptome changes.

For over a decade, myosin VI has been linked to transcription (Cook et al., 2018, Fili et al., 2020, Fili et al., 2017, Große-Berkenbusch et al., 2020, Vreugde et al., 2006) and here we have gained further understanding of its nuclear function, including its interaction with nuclear receptors and DNA (Fili et al., 2017). Furthermore, until now, it has not been possible to determine the precise role that myosin VI plays in this vital process and why the properties of a myosin would be required for transcription. As with many nuclear proteins (Carmo-Fonseca, 2002, Cook, 2010, Cremer et al., 2006, Verschure et al., 1999), we have shown that myosin VI forms molecular clusters, possibly to enhance its activity and the efficiency of biochemical processes. The formation of these clusters is ATP and actin-dependent and, therefore, relies on the motor properties of myosin VI. Here, we have dissected the molecular mechanistic need for myosin VI to be capable of switching from a transporter to an anchor in a force induced manner. Therefore, we have been able to directly address why a nuclear myosin, with its biophysical properties to sense and respond to force, is required in transcription in order to hold RNAPII *in situ*.

Overall, we present a model, whereby myosin VI anchors RNAPII at, or near, sites of transcription initiation (Figure 9) within the nucleus. This approach could enable enhanced RNAPII binding to initiate transcription and facilitate rapid recycling of RNAPII to drive higher expression levels.

**Figure 9.**
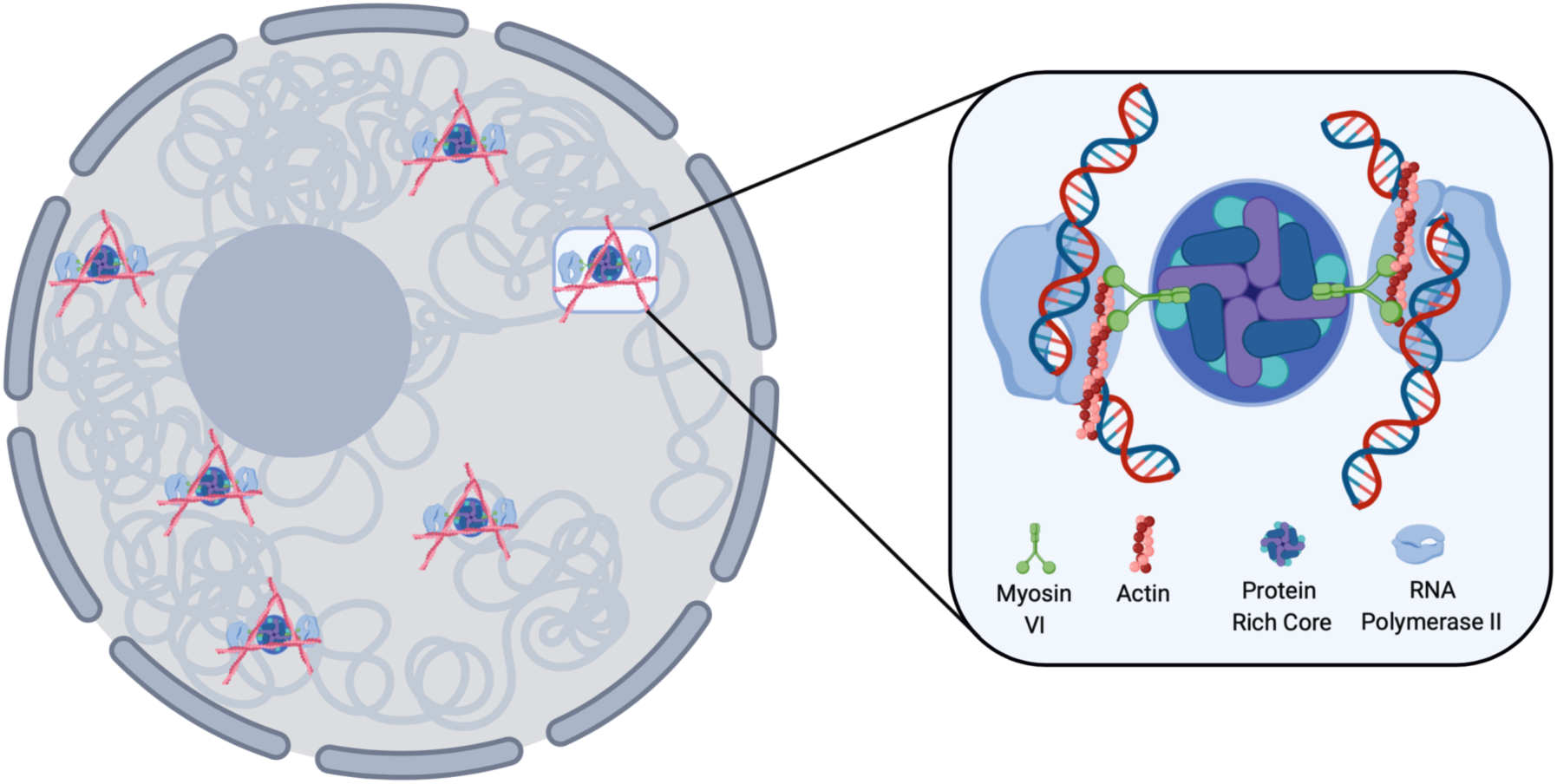
Schematic of myosin VI anchoring RNAPII at transcription factories. Based on the data presented here we can propose this model for RNAPII clustering. Transcription sites within the nucleus are marked by clustering of RNAPII and actin filaments. Focusing on the clusters: The protein rich core consists of transcription regulators which the DNA would also contact. Myosin VI interacts with these proteins and DNA via its C-terminal domain. Myosin VI then interacts with RNAPII through nuclear actin filaments thereby providing a framework for mechanical linkage to RNAPII.

Such a mechanism would also fit with the observation that myosin VI functions in gene pairing (Zorca et al., 2015). We have previously shown that myosin VI interacts with RNAPII through actin (Fili et al., 2017), present within the RNAPII complex. Given that the interaction of myosin VI with nuclear receptors and DNA is mediated by its CBD, we propose that myosin VI could be bound to DNA and/or transcription regulators via the CBD, whilst simultaneously interacting with RNAPII through actin. We propose that the myosin VI orientation is critical to enable force-induced anchoring of RNAPII. These results are consisted with recent reports of nuclear actin clustering during transcription stimulation (Wei et al., 2020). Interestingly, myosin VI is the only actin minus-end motor protein, therefore actin polymerizing from these sites of transcription would also provide a framework to recruit myosin VI to these sites and subsequently to RNAPII.

It would be interesting to address whether RNAPII moves along the DNA, or whether DNA is trafficked through the factories. Published ChIP data (Vreugde et al., 2006) showed that myosin VI is present throughout the gene body. Based on the clustering behaviour of myosin VI and our model, we would expect DNA to be trafficked through these static factories. Maintaining the interaction would support rapid recycling of RNAPII, as mentioned above.

Nuclear Myosin I, has a well-established role in transcription and this family of myosins is also capable of acting as force-induced anchors (Greenberg and Ostap, 2013). It remains to be explored if both proteins interact with RNAPII simultaneously to stabilise the complex or if, conversely, the different myosin proteins interact with distinct populations of RNAPII. Overall, it would not be surprising if nuclear myosins are deployed in a similar way in other nuclear processes such as DNA repair, where myosin proteins are also known to function (Caridi et al., 2018, Kulashreshtha et al., 2016, Venit et al., 2020).

Upon perturbation of myosin VI, RNAPII was observed around the nuclear periphery, which highlights the key role of myosin VI in maintaining the nuclear organisation of RNAPII. The basis for this localisation is not yet clear, but we can postulate two mechanisms. The protein may be en-route for nuclear export as part of a degradation process which initiates once transcription is disrupted (Muratani and Tansey, 2003). Alternatively, RNAPII may be excluded from the chromatin body as DNA condensation takes place, through the increase of repressive transcription histone markers. This would assume that the loss of transcription triggers a default to a repressive marker. In this manner, the now highly dynamic RNAPII would move into less dense chromatin regions, which may speculatively be present at the periphery. Moreover, an increase in chromatin condensation would further increase RNAPII dynamics, by reducing RNAPII interacts with DNA and lead to an increase in non-specific interactions. In our model, myosin VI anchoring would also lead to a reduction in these non-specific interactions by holding the complex at the active regions.

The ability of RNAPII to cluster has been shown to be important for transcription (Cho et al., 2016a, Cho et al., 2016b). Our findings further support this, given the disruptive effect of myosin VI perturbation on the organisation of RNAPII and, subsequently, on transcription. The simultaneous expression of genes from a single transcription factory ensures that numerous proteins required for a specific pathway are efficiently produced in a co-ordinated fashion. Clustering of RNAPII has been shown to be dynamic with structures lasting around 10 seconds (Cho et al., 2016a, Cho et al., 2016b) and myosin VI has also been shown to display dynamic binding (up to 20 seconds) (Große-Berkenbusch et al., 2020). We therefore conclude that there is likely to be turnover of proteins within these clusters, however we suggest that myosin VI enhances the RNAPII binding time to enable transcription initiation. Phase separation drives the formation of membraneless compartments through cooperative interactions between molecules (Hnisz et al., 2017). This process has been shown to contribute to the regulation of transcription, with impacts upon enhancers, mediators and the RNAPII C-Terminal-Domain (Boehning et al., 2018, Cho et al., 2018, Sabari et al., 2018). In contrast, our results suggest there are underlying mechanical processes contributing to the establishment of transcription factories with regard to RNAPII. Importantly, phase separation observations are still consistent with our conclusions because these processes can occur locally within the factories and may contribute to larger genomic rearrangements by bringing chromatin to the factories.

The formation of transcription factories has been proposed to be linked to transcription stimulation events. According to our model, we expect the need for myosins to be deployed in conditions of high transcriptional load to increase activate and/or recycling of RNAPII. Indeed, our RNA-seq analysis suggests that myosin VI plays a key role during stimulation and the downstream cell response processes. We therefore propose that the function of myosin VI is particularly critical when cells are under high transcription load and undergo rapid changes in the gene expression landscape. This is consistent with previous observations of myosin VI being over-expressed in several cancers, along with interacting with nuclear receptors and participating in the expression of target genes (Dunn et al., 2006, Loikkanen et al., 2009, Puri et al., 2010, Wang et al., 2016, Wang et al., 2015, Wollscheid et al., 2016).

With regard to the factories, various key questions remain unanswered, for instance how the clustering sites are selected and how they are brought together? What is the internal structure of the clusters with regard to positioning of myosin VI, RNAPII, DNA and transcription factors? Although processes such as chromatin condensation and genome organisation such as Topologically Associating Domains, could play a role in dictating where the clustering sites form, the underlying mechanism remains elusive. Myosin VI has been shown to interact with nuclear receptors therefore it could be that myosin VI is recruited to these binding sites and clusters subsequently build around these locations. Loop exclusion may then have the ability to cluster local sites together (Schoenfelder and Fraser, 2019). Moreover, as it has been proposed (Große-Berkenbusch et al., 2020), we could speculate that the cargo transportation ability of nuclear myosins, including myosin VI, could be harnessed in combination with actin filaments, in order to bring genes together across large distances. Understanding the internal organisation of the clusters would require high-resolution structure characterisation in the cell and with reconstituted complexes.

In summary, we have investigated the function of nuclear myosin VI and uncovered a fundamental role in the spatial organisation of gene expression. This appears to be critical for transcription stimulation events where multiple genes are simultaneously expressed and cells rapidly adapt to environmental changes.

## ACKNOWLEDGEMENTS

We thank the UKRI-MRC (MR/M020606/1), UKRI-STFC (19130001) and the Royal Society (IES\R3\183138) for funding to C.P.T. ERC StG 637987 and SPP 2202 GE 2631/1-1 for funding to J.C.M.G. Aberration-corrected multi-focal microscopy was performed in collaboration with the Advanced Imaging Center at Janelia Research Campus, a facility jointly supported by the Howard Hughes Medical Institute and the Gordon and Betty Moore Foundation. We also thank Darren Griffin (University of Kent) and Alessia Buscaino (University of Kent) for sharing of equipment, and Satya Khuon (Janelia Research Campus for assisting with cell culture. JF549 dyes were kindly provided by Luke Lavis (Janelia Research Campus). We also thank grant from NYU Abu Dhabi, The Swedish Research Council and Cancerfonden to P.P. and we acknowledge technical help from the NYU Abu Dhabi Center for Genomics and Systems Biology, in particular Marc Arnoux and Mehar Sultana. We appreciate the computational platform provided by NYUAD HPC team and are especially thankful to Nizar Drou for technical help.

## AUTHOR CONTRIBUTIONS

C.P.T. conceived the study. Y.H-G., N.F., A.dS., R.E.G., A.W.C. and C.P.T. designed experiments. N.F., Y.H-G. and C.P.T. designed and cloned constructs. Y.H-G., A.G-B. and J.C.M.G. created cell lines. Y.H-G., N.F., A.dS. and C.P.T. performed single molecule imaging experiments. Imaging was supported by L.W., M.M-F. and J.A. Y.H-G., A.W.C., T.V., and P.P. performed and analyzed the genomics experiments. N.F., R.E.G., H.C.W.R. and C.P.T. expressed, purified and performed experiments with recombinant proteins. L.W., J.A., E.W., T-L.C., A.dS. and C.P.T. contributed to single molecule data analysis. C.P.T. supervised the study. C.P.T. wrote the manuscript with comments from all authors.

## EXPERIMENTAL PROCEDURES

### Constructs

A list of constructs are provided in Supplementary Table 2. Constructs generated in this work are described below. Halo or SNAP tags were used through to provide a specific protein labeling strategy for live cells (Toseland, 2013). The SNAP-Rpb1 construct was generated by sub-cloning the SNAP tag from the pSNAP_f_-C1 plasmid (Addgene 58186) into the NheI and SacII of the pHalo-Rbp1 plasmid (A gift from Darzacq lab), following removal of the Halo tag. pcDNA3.1 Halo-MVI, pcDNA3.1 Halo-MVI-Spring, pFastbac Halo-MVI-Spring, and pFastbac Halo-MVI-Spring bZip were ordered as synthetic constructs.

### Protein Expression using Baculovirus system

Full-length MVI NI, MVI Spring, MVI Spring bZip (1-914) and *Xenopus* calmodulin were expressed in *Sf9* and *Sf21* (*Spodoptera frugiperda*) insect cells using the Bac-to-Bac® Baculovirus Expression System (Invitrogen). *Sf9* cells were cultured in Sf900 media (Gibco). Recombinant bacmids were generated following the manufacturer’s instructions and were transfected into adherent Sf9 cells to generate the P1 viral stock. *Sf9* cells were infected in suspension at 27ºC and 100 rpm with 1 in 50 dilution of P1 and P2 viral stocks to yield P2 and P3 stocks, respectively. Finally, expression of recombinant proteins was set up by infecting *sf21* cells with the P3 viral stock in Spodopan media (PAN Biotech). To ensure correct folding of the myosin VI constructs, cells were simultaneously infected with P3 viral stock of the myosin VI constructs together with calmodulin at a 0.75 ratio The cells were harvested after 3 days by centrifugation for 15 min at 700xg and at 4 °C and resuspended in ice cold myosin extraction buffer (90 mM KH_2_PO_4_, 60 mM K_2_HPO_4_, 300 mM KCl, pH 6.8), supplemented with Proteoloc protease inhibitor cocktail (Expedeon) and 100 µM PMSF, before proceeding to protein purification. Prior to sonication, an additional 5 mg recombinant calmodulin was added together with 2 mM DTT. After sonication, 5 mM ATP and 10 mM MgCl_2_ were added and the solution was rotated at 4 °C for 30 min before centrifugation (20,000g, 4°C, 30 min). Then, the cell lysate was subjected to the purification. Proteins were purified by affinity chromatography (HisTrap FF, GE Healthcare). The purest fractions were further purified through a Superdex 200 16/600 column (GE Healthcare).

### Cell culture and Transfection

HeLa (ECACC 93021013) cells were cultured at 37ºC and 5% CO_2_, in Gibco MEM Alpha medium with GlutaMAX (no nucleosides), supplemented with 10% Fetal Bovine Serum (Gibco), 100 units/ml penicillin and 100 µg/ml streptomycin (Gibco). For the transient expression of myosin VI and mutants, HeLa cells grown on glass coverslips were transfected using Lipofectamine 2000 (Invitrogen), following the manufacturer’s instructions. Depending on the construct, 24 h – 72 h after transfection, cells were subjected to nuclear staining using Hoechst 33342 (Thermo Scientific), fixed and analysed or subjected to indirect immunofluorescence (see below).

### Cell Treatments

For MVI knock-down experiments, HeLa cell monolayers, seeded to 30 – 50 % confluency, were transfected with human myosin VI siRNA duplex (5′GGUUUAGGUGUUAAUGAAGtt-3′) (Ambion) or AllStars Negative Control siRNA duplex (Qiagen) at a concentration of 50 nM, using Lipofectamine 2000 (Invitrogen), according to the manufacturer’s guidelines. Cells were fixed or harvested after 48 h for further analysis. To inhibit myosin VI, cells were treated with 25 µM TIP (Sigma) for 1 h at 37 ºC. To inhibit actin polymerization, cells were treated with 1 µM Latrunculin B (Sigma) for 1h at 37 ºC. To inhibit RNAPII transcription, cells were treated with 5 µg/ml α-amanitin (Sigma) for 4h at 37 ºC. For serum stimulation, 4.8×10^5^ Hela cells were seeded in DMEM complete media in 6 well plates to achieve 70-80% confluency on the following day. For serum starvation, cells were grown in DMEM with 0.5% FBS at 37 ºC for 24 h. To stimulate the starved cells, media was replaced with complete media containing 10% FBS for another 24 h after which cells were fixed for immunofluorescence.

### Stable cell line generation

The stable cell lines used in this study are named as HeLa-Halo MVI (pHalo-MVI vector stably expressed in HeLa) and Hela-Halo Rpb1 (pHalo-Rpb1 vector stably expressed in HeLa). The Hela-Halo MVI were generated as described in (Große-Berkenbusch et al., 2020). To generate Hela cells stably expressing pHalo-Rpb1, the plasmid was transfected in 6-well plates using lipofectamine 2000 protocol (Thermo Fisher Scientific). The transfected cells were selected using optimal concentrations of G418 antibiotic (G418 Sulfate, Gibco) in the complete media (0.5mg/ml) for 9-10 days until most of the untransfected cells were dead and those survived would have integrated the desired plasmid. The cells were harvested when they reached about 60-70% confluency and were expanded into multiple T75 flasks with 1:10 ratio. Some cells at this stage were seeded onto coverslips and stable transfection of desired plasmids was confirmed by using specific fluorescent ligands to Halo-tag (TMR, Promega). The cells seeded in T75 flasks were allowed to grow for further 3-4 weeks in complete media with G418 replaced twice a week. When the cells reached high confluency, they were frozen down as polyclonal stable cell line.

### Immunofluorescence

Transfected and non-transfected HeLa cells were fixed for 15 min at room temperature in 4% (w/v) paraformaldehyde (PFA) in PBS and residual PFA was quenched for 15 min with 50 mM ammonium chloride in PBS. All subsequent steps were performed at room temperature. Cells were permeabilised and simultaneously blocked for 15 min with 0.1 % (v/v) Triton X-100 and 2 % (w/v) BSA in PBS. Cells were then immuno-stained against the endogenous proteins by 1 h incubation with the indicated primary and subsequently the appropriate fluorophore-conjugated secondary antibody (details below), both diluted in 2 % (w/v) BSA in PBS. When using anti-phospho antibodies, immunofluorescence protocol was performed in TBS. The following antibodies were used at the indicated dilutions: Rabbit anti-myosin VI (1:200, Atlas-Sigma HPA0354863), Rabbit anti-Histone H3 (tri methyl K9) (1:500, Abcam ab8898), Rabbit anti-Histone H3 (acetyl K27) (1:500, Abcam ab4729), Rabbit anti-Histone H3 (acetyl K9) (1:200, Abcam ab4441), Rabbit anti-RNAPII phospho Ser5 (1:500, Abcam Ab5131), Mouse anti-RNAPII phospho Ser5 (1:500, Abcam Ab5408), Donkey anti-mouse Alexa Fluor 488-conjugated (1:500, Abcam Ab181289), Donkey anti-rabbit Alexa Fluor 647-conjugated (1:500, Abcam Ab181347) and Donkey anti-rabbit Alexa Fluor 488-conjugated antibody (1:500, Abcam Ab181346). Coverslips were mounted on microscope slides with Mowiol (10% (w/v) Mowiol 4-88, 25% (w/v) glycerol, 0.2 M Tris-HCl, pH 8.5), supplemented with 2.5% (w/v) of the anti-fading reagent DABCO (Sigma).

### Immunoblot Analysis

The total protein concentration was determined by Bradford Assay (Sigma) following the manufacturer’s instructions. Cell lysates were heat-denatured and resolved by SDS-PAGE. The membrane was probed against the endogenous proteins by incubation with primary Rabbit anti-myosin VI (1:500, Atlas-Sigma HPA0354863-100UL) or Mouse anti-RNAPII phospho Ser5 (1:500, Abcam Ab5408) and subsequently secondary Goat anti-rabbit antibody (1:15000 Abcam ab6721) or Goat anti-mouse antibody (1:15000, Abcam ab97023) coupled to horseradish peroxidase. The bands were visualised using the ECL Western Blotting Detection Reagents (Invitrogen) and the images were taken using Syngene GBox system. Images were processed in ImageJ.

### Fluorescence Imaging

Cells were visualised using either the ZEISS LSM 880 confocal microscope or the widefield Olympus IX71 microscope. The former was equipped with a Plan-Apochromat 63x 1.4 NA oil immersion lens (Carl Zeiss, 420782-9900-000). Three laser lines, i.e. 405 nm, 488 nm and 561 nm, were used to excite the fluorophores, i.e. Hoechst, GFP and RFP, respectively. The built-in dichroic mirrors (Carl Zeiss, MBS-405, MBS-488 and MBS-561) were used to reflect the excitation laser beams on to cell samples. The emission spectral bands for fluorescence collection were 410 nm-524 nm (Hoechst), 493 nm-578 nm (GFP) and 564 nm-697 nm (RFP). The detectors consisted of two multi anode photomultiplier tubes (MA-PMT) and 1 gallium arsenide phosphide (GaAsP) detector. The green channel (GFP) was imaged using GaAsP detector, while the blue (Hoechst) and red (RFP) channels were imaged using MA-PMTs. ZEN software (Carl Zeiss, ZEN 2.3) was used to acquire and render the confocal images. The later was equipped with an PlanApo 100xOTIRFM-SP 1.49 NA lens mounted on a PIFOC z-axis focus drive (Physik Instrumente, Karlsruhe, Germany), and illuminated with an automated 300W Xenon light source (Sutter, Novato, CA) with appropriate filters (Chroma, Bellows Falls, VT). Images were acquired using a QuantEM (Photometrics) EMCCD camera, controlled by the Metamorph software (Molecular Devices). The whole volume of cells was imaged by acquiring images at z-steps of 200 nm. Widefield images were deconvolved with the Huygens Essential version 17.10 software. Confocal Images were deconvolved using the Zeiss Zen2.3 Blue software, using the regularised inverse filter method. All images were then analysed by ImageJ.

### STORM Imaging

Cells were seeded on pre-cleaned No 1.5, 25-mm round glass coverslips, placed in 6-well cell culture dishes. Glass coverslips were cleaned by incubating them for 3 hours, in etch solution, made of 5:1:1 ratio of H_2_O: H_2_O_2_ (50 wt. % in H_2_O, stabilized, Fisher Scientific): NH_4_OH (ACS reagent, 28-30% NH_3_ basis, Sigma), placed in a 70°C water bath. Cleaned coverslips were repeatedly washed in filtered water and then ethanol, dried and used for cell seeding. Transfected or non-transfected cells were fixed in pre-warmed 4% (w/v) PFA in PBS and residual PFA was quenched for 15 min with 50 mM ammonium chloride in PBS. Immunofluorescence (IF) was performed in filtered sterilised PBS, unless when anti-phospho antibodies were used. Then, IF was performed in filtered sterilised TBS. Cells were permeabilized and simultaneously blocked for 30 min with 3% (w/v) BSA in PBS or TBS, supplemented with 0.1 % (v/v) Triton X-100. Permeabilized cells were incubated for 1h with the primary antibody and subsequently the appropriate fluorophore-conjugated secondary antibody, at the desired dilution in 3% (w/v) BSA, 0.1% (v/v) Triton X-100 in PBS or TBS. The antibody dilutions used were the same as for the normal IF protocol (see above), except from the secondary antibodies which were used at 1:250 dilution. Following incubation with both primary and secondary antibodies, cells were washed 3 times, for 10 min per wash, with 0.2% (w/v) BSA, 0.05% (v/v) Triton X-100 in PBS or TBS. Cells were further washed in PBS and fixed for a second time with pre-warmed 4% (w/v) PFA in PBS for 10 min. Cells were washed in PBS and stored at 4 °C, in the dark, in 0.02% NaN3 in PBS, before proceeding to STORM imaging.

Before imaging, coverslips were assembled into the Attofluor® cell chambers (Invitrogen). Imaging was performed in freshly made STORM buffer consisting of 10 % (w/v) glucose, 10 mM NaCl, 50 mM Tris – pH 8.0, supplemented with 0.1 % (v/v) 2-mercaptoethanol and 0.1 % (v/v) pre-made GLOX solution which was stored at 4 0C for up to a week (5.6 % (w/v) glucose oxidase and 3.4 mg/ml catalase in 50 mM NaCl, 10 mM Tris – pH 8.0). All chemicals were purchased from Sigma. Imaging was undertaken using the Zeiss Elyra PS.1 system. Illumination was from a HR Diode 642 nm (150 mW) and HR Diode 488 nm (100 mW) lasers where power density on the sample was 7-14 kW/cm^2^ and 7-12 kW/cm^2^, respectively Imaging was performed under highly inclined and laminated optical (HILO) illumination to reduce the background fluorescence with a 100x NA 1.46 oil immersion objective lens (Zeiss alpha Plan-Apochromat) with a BP 420-480/BP495-550/LP 650 filter. The final image was projected on an Andor iXon EMCCD camera with 25 msec exposure for 20000 frames.

Image processing was performed using the Zeiss Zen software. Where required, two channel images were aligned following a calibration using a calibration using pre-mounted MultiSpec bead sample (Carl Zeiss, 2076-515). For calibration, a 2 µm Z-stack was acquired at 100 nm steps. The channel alignment was then performed in the Zeiss Zen software using the Affine method to account for lateral, tilting and stretching between the channels. The calibration was performed during each day of measurements.

The images were then processed through our STORM analysis pipeline using the Zen software. Single molecule detection and localisation was performed using a 9 pixel mask with a signal to noise ratio of 6 in the “Peak finder” settings while applying the “Account for overlap” function. This function allows multi-object fitting to localise molecules within a dense environment. Molecules were then localised by fitting to a 2D Gaussian.

The render was then subjected to model-based cross-correlation drift correction and detection grouping to remove detections within multiple frames. Typical localisation precision was 20 nm for Alexa-Fluor 647 and 30 nm for Alexa-Fluor 488. The final render was then generated at 10 nm/pixel and displayed in Gauss mode where each localisation is presented as a 2D gaussian with a standard deviation based on its precision. The localisation table was exported as a csv for import in to Clus-DoC.

### Clus-DoC

The single molecule positions were exported from Zeiss Zen Black version and imported into the Clus-DoC analysis software (Pageon et al., 2016) (https://github.com/PRNicovich/ClusDoC). The region of interest was determined by the nuclear staining. First the Ripley K function was completed on each channel identifying the r max. The r max was then assigned for DBSCAN if one channel was being analysed or Clus-Doc if two channel colcalisation was being analysed. The clustering size was set to a minimum of 5 molecules within a cluster with a smoothing of a cluster being set at 7 nm. All other analyses parameters remained at default settings. Data concerning each clusters was exported and graphed using Plots of Data.

### High content imaging

Cells were seeded onto Corning® 384 well microplates at a density of 5,000 cells per well. The cells were grown for 24 hours, followed by the necessary treatments. The cells were fixed and immunofluorescence was undertaken as described above, due to the cells being grown directly on the plates no mounting of coverslips was required. Stained cells in plate were scanned via Cellomics ArrayScan™ XTI High Content Analysis (HCS) platform (Thermo Fisher Scientific), with a 20x Objective. Compartment Analysis Bio Application software (Cellomics) was applied to quantitatively analyse the immunostaining spots in the nucleus based on a mask created using the nuclear Hoechst staining. For each experiment, at least 1000 valid single cells per culture well were quantified and at least 10 independent culture wells (10 biological replicates) were analysed, fluorescence intensities were then plotted using Prism 8, Graphpad.

### Size-exclusion Chromatography

100 µl samples of 2mg/ml purified protein, was applied to a Superdex 200 (30 × 1 cm) analytical column (GE Healthcare) equilibrated in 150 mM NaCl, 50 mM Tris.HCl (pH 7.5) and 1 mM DTT and controlled using Waters 626 HPLC and OMNISEC (Malvern Panalytical) at room temperature.

### Multi-focal Imaging and Particle Tracking Analysis

Cells stably or transiently expressing Halo-tag or SNAP-tag constructs were labelled for 15 min with HaloTag-JF549 or SNAP-tag-JF549 ligand, respectively, in cell culture medium at 37°C, 5% CO_2_. 10 nM ligand was used to label Halo-tagged myosin VI constructs, whereas 50 nM ligand was used to label Halo- or SNAP-tagged RNAPII. Cells were washed for 3 times with warm cell culture medium and then incubated for further 30 min at 37°C, 5% CO_2_. Cells were then washed three times in pre-warmed FluoroBrite DMEM imaging medium (ThermoFisher Scientific), before proceeding to imaging.

Single molecule imaging was performed using an aberration-corrected multifocal microscope (acMFM), as described by Abrahamsson et al. (Abrahamsson et al., 2013). Briefly, samples were imaged using 561nm laser excitation, with typical irradiance of 4-6 kW/cm^2^ at the back aperture of a Nikon 100x 1.4 NA objective. Images were relayed through a custom optical system appended to the detection path of a Nikon Ti microscope with focus stabilization. The acMFM detection path includes a diffractive multifocal grating in a conjugate pupil plane, a chromatic correction grating to reverse the effects of spectral dispersion, and a nine-faceted prism, followed by a final imaging lens.

The acMFM produces nine simultaneous, separated images, each representing successive focal planes in the sample, with ca. 20 µm field of view and nominal axial separation of ca. 400nm between them. The nine-image array is digitized via an electron multiplying charge coupled device (EMCCD) camera (iXon Du897, Andor) at up to 32ms temporal resolution, with typical durations of 30 seconds.

3D+t images of single molecules were reconstructed via a calibration procedure, implemented in Matlab (MathWorks), that calculates and accounts for (1) the inter-plane spacing, (2) affine transformation to correctly align each focal plane in the xy plane with respect to each other, and (3) slight variations in detection efficiency in each plane, typically less than ±5-15% from the mean.

Reconstructed data were then subject to pre-processing, including background subtraction, mild deconvolution (3-5 Richardson-Lucy iterations), and/or Gaussian de-noising prior to 3D particle tracking using the MOSAIC software suite (Sbalzarini and Koumoutsakos, 2005). Parameters were set where maximum particle displacement was 400 nm and a minimum of 10 frames was required. Tracks were reconstructed, and diffusion constants were extracted via MSD analysis (Aaron et al., 2019) using custom Matlab software assuming an anomalous diffusion model.

### Circular dichroism Spectroscopy

1 mgmL^−1^ of protein was analysed using far UV spectra (190nm-270nm) measured by a Jasco J715 Circular Dichroism Spectrometer (Jasco Inc.). Spectra were taken at 20°C. 4 readings were taken for each measurement and averaged by the software provided. For spectra analysis the following equation was used.

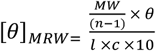

Where *θ_MRW_* is the mean residue elipticity, *MW* is the molecular weight of the protein, *n* is the number of amino acids, *θ* is the degrees in elipticity, *l* is the path length and *c* is the concentration.

### Incucyte live cell imaging

Cells were seeded onto 96-well tissue culture dishes at equal densities in 6 replicates. After attachment overnight, cells were transfected with MVI siRNA, or scrambled siRNA (Qiagen). Photomicrographs were taken every hour using an IncuCyte live cell imager (Essen Biosciences, Ann Harbor, MI) and confluency of cultures was measured using IncuCyte software. Confluency values between wells were normalised to initial confluency for comparison.

### RNA extraction and RT-qPCR

RNA from HeLa cells was extracted using Gene Jet RNA purification kit (Thermo scientific) according to manufacturer’s protocol. The RNA concentration was measured using Geneflow Nanophotometer and RT-qPCR was performed with one-step QuantiFast SYBR Green qPCR kit (Qiagen) using 50ng of RNA in each sample. A list of qPCR primers is given in Supplementary Table 3.

### RNA-seq and analysis

Total RNA was extracted from three replicates of WT, MVI KD and Scrambled siRNA. Ice cold TRIzol reagent was added to each culture and homogenised. The mixture was then incubated for 5 mins at room temperature then chloroform was added to the lysis and incubated for 3 mins. The samples were then centrifuged at 12,000 × g at 4°C. The colourless aqueous phase was collected. The RNA was then precipitated with incubation for 10 mins with isopropanol before centrifugation for a further 10 mins at 12,000 × g at 4 °C. The pellet was washed in 75% (v/v) ethanol, vortexed and centrifuged for 5 mins at 7500 × g at 4°C. The RNA pellet is air dried for 10 mins. The pellet is then resuspended in 50µL of RNase-free water containing 0.1mM EDTA and incubated at 55°C for 15 mins to allow the RNA to dissolve. The RNA was then quantified using then 260nm absorbance, ensuring the A260/A280 ratio was approximately 2, therefore implying the sample is pure. The sample was then further purified using the RNeasy kit (Qiagen) where the manufacturers protocol was followed exactly. Once the purity and stability had been measured the RNA was then stored at −80°C.

The RNA-seq libraries were prepared with TruSeq RNA Library Prep kit v2 as per protocol instructions. Resulting libraries concentration, size distribution and quality were assessed on a Qubit fluorometer with a dsDNA high sensitivity kit and on an Agilent 2100 bioanalyzer using a DNA 7500 kit. Then libraries were normalized, pooled and quantified with a KAPA Library quantification kit for Illumina platforms on a ABI StepOnePlus qPCR machine, then loaded on a high output flow cell and paired-end sequenced (2×75 bp) on an Illumina NextSeq 550 next generation sequencer (performed at the NYUAD Sequencing Center). The raw FASTQ reads were quality trimmed using Trimmomatic (version 0.36) (Bolger et al., 2014) to trim low quality bases, systematic base calling errors, as well sequencing adapter contamination. FastQC (www.bioinformatics.babraham.ac.uk/projects/fastqc) was used to assess the quality of the sequenced reads pre/post quality trimming. Only the reads that passed quality trimming in pairs were retained for downstream analysis. The quality trimmed RNAseq reads were aligned to the Homo sapiens GRch38.p4 genome using HISAT2 (version 2.0.4) (Kim et al., 2015). The resulting SAM alignment files were then converted to BAM format and sorted by coordinate using SAMtools (version 0.1.19) (Li et al., 2009). The BAM alignment files were processed using HTseq-count (Anders et al., 2015) using the reference annotation file to produce raw counts for each sample. The raw counts were then analyzed using the online analysis portal NASQAR (http://nasqar.abudhabi.nyu.edu/) in order to merge, normalize and identify differentially expressed genes by using the START app (Nelson et al., 2017). Differentially expressed genes by at least 2-fold log2(FC)≥1 and adjusted p-value of <0.05 for upregulated genes and log2(FC)≤-1 and adjusted p-value of <0.05 for downregulated genes) between the samples which were then subjected to Gene Ontology (GO) enrichment using ShinyGo v0.60 (http://bioinformatics.sdstate.edu/go/) (Ge et al., 2019).

### Chromatin Immunoprecipitation (ChIP)

To identify specific RNAPII-DNA interactions, ChIP was performed using mouse anti-RNAPII-pSer5 antibody (Abcam Ab5408). A confluent T175 flask (10×10^6^ – 30×10^6^) of HeLa cells was crosslinked by adding formaldehyde dropwise directly to the media to a final concentration of 0.75% and was left for gentle rotation at room temperature (RT) for 10min. To stop the reaction, glycine was added to a final concentration of 125 mM and was incubated with shaking for 5 min at RT. The cells were washed twice with 10 ml of cold PBS and were scraped in 5-8 ml of cold PBS. All cells were collected and centrifuged at 1000xg, 4°C for 5 min. The pellet was re-suspended in ChIP lysis buffer (750 μL per 1×10^7^ cells) (lysis buffer: 50 mM HEPES-KOH pH 7.5, 140 mM NaCl, 1 mM EDTA pH8, 1 % TritonX-100, 0.1 % Sodium Deoxycholate, 0.1 % SDS and Protease Inhibitors) and was incubated on ice for 10 min. The cells were sonicated using the diagenode bioruptor sonicator in order to shear DNA to an average fragment size of 200-800 bp. The fragment size was analysed on a 1.5 % agarose gel. After sonication, cell debris was removed by centrifugation for 10 min, 4°C, 8000xg and the supernatant (chromatin) was used for the immunoprecipitation. The sonicated chromatin was snap frozen on dry ice and was stored at −80°C until further use (max storage 3 months).

The chromatin prepared above was diluted 1:10 with RIPA buffer (50 mM Tris-HCl pH8, 150 mM NaCl, 2 mM EDTA pH8, 1% NP-40, 0.5% Sodium Deoxycholate, 0.1% SDS and Protease Inhibitors) and was distributed into 6 tubes (approximately 1×10^6^ cells per IP) – 3 samples for specific antibody (MVI) and 3 samples for the no antibody control (beads only). 10% of diluted chromatin was removed to serve as input sample and was stored at −20°C until further use. All chromatin samples were pre-cleared using the protein A magnetic beads (Thermo Fisher Scientific) for 30 min after which 20 µl of mouse anti-RNAPII-pSer5 antibody was added to each of the triplicate Ab samples (1 in 50 dilution) and the tubes were rotated at 4°C, overnight. Next day, 40 µl of protein A magnetic beads (washed three times in RIPA buffer) were added to each of the samples including the no antibody control tubes and were put on rotation at 4°C for 1 h. After 1h, the beads were collected using a magnetic rack and were washed twice in low salt buffer (0.1 % SDS, 1 % Triton X-100, 2 mM EDTA, 20 mM Tris-HCl pH 8, 150 mM NaCl), once in high salt buffer (0.1 % SDS, 1 % Triton X-100, 2 mM EDTA, 20 mM Tris-HCl pH 8, 500 mM NaCl), once in LiCl wash buffer (0.25 M LiCl, 1 % NP-40, 1 % Sodium Deoxycholate, 1 mM EDTA, 10 mM Tris-HCl pH 8) and finally in TE (10mM Tris pH8, 1mM EDTA). DNA was eluted by adding 120 µl of elution buffer (1 % SDS, 100 mM NaHCO_3_) to the beads and vortexing them slowly for 15 min at 30°C. To reverse crosslink the protein-DNA complexes, 4.8 µl of 5M NaCL and 2 µl RNase A (10mg/ml) was added to the elutes including the input sample that was stored at −20°C and they were incubated while shaking at 65°C overnight followed by proteinase K treatment at 60°C for 1 h. The DNA was then purified using phenol:chloroform extraction and the samples were analysed by qPCR using primers in Supplementary Table 3.

### Steady-state ATPase Activity

Ca^2+^-actin monomers were converted to Mg^2+^-actin with 0.2 mM EGTA and 50 µM MgCl_2_ before polymerizing by dialysis into 20 mM Tris.HCl (pH7.5), 20 mM imidazole (pH 7.4), 25 mM NaCl and 1 mM DTT. A 1.1 molar equivalent of phalloidin (Sigma) was used to stabilize actin filaments (Batters et al., 2012, Toseland, 2014).

Steady-state ATPase activities were measured at 25 °C in KMg50 buffer (50 mM KCl, 1 mM MgCl_2_, 1 mM EGTA, 1 mM DTT, and 10 mM imidazole, pH 7.0). Supplemented with the NADH-coupled assay components, 0.2 mM NADH, 2 mM phosphoenolpyruvate, 3.3 U ml^−1^ lactate dehydrogenase, 2.3 U ml^−1^ pyruvate kinase and various actin concentrations (0 – 30 µM). The final [Mg.ATP] was 5 mM and MVI concentration was 100– 300 nM. The assay was started by the addition of MVI. The change in absorption at OD_340_ nm was followed for 5 min. The *k*_cat_ and *K*actin values were determined by fitting the data to equation:

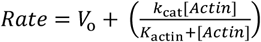

*V*o is the basal ATPase activity of MVI, *k*_cat_ is the maximum actin-activated ATPase rate and *K*_actin_ is the concentration of actin needed to reach half maximal ATPase activity.

### Graphics

Unless stated, data fitting and plotting was performed using Plots of data (Postma and Goedhart, 2019) and Grafit Version 5 (Erithacus Software Ltd). Cartoons were generated using the BioRender software (www.biorender.com).

### Data Availability

The data supporting the findings of this study are available from the corresponding author on request.

