## Supplementary Information and Figures for "Nuclear myosin VI regulates the spatial organization of mammalian transcription initiation"

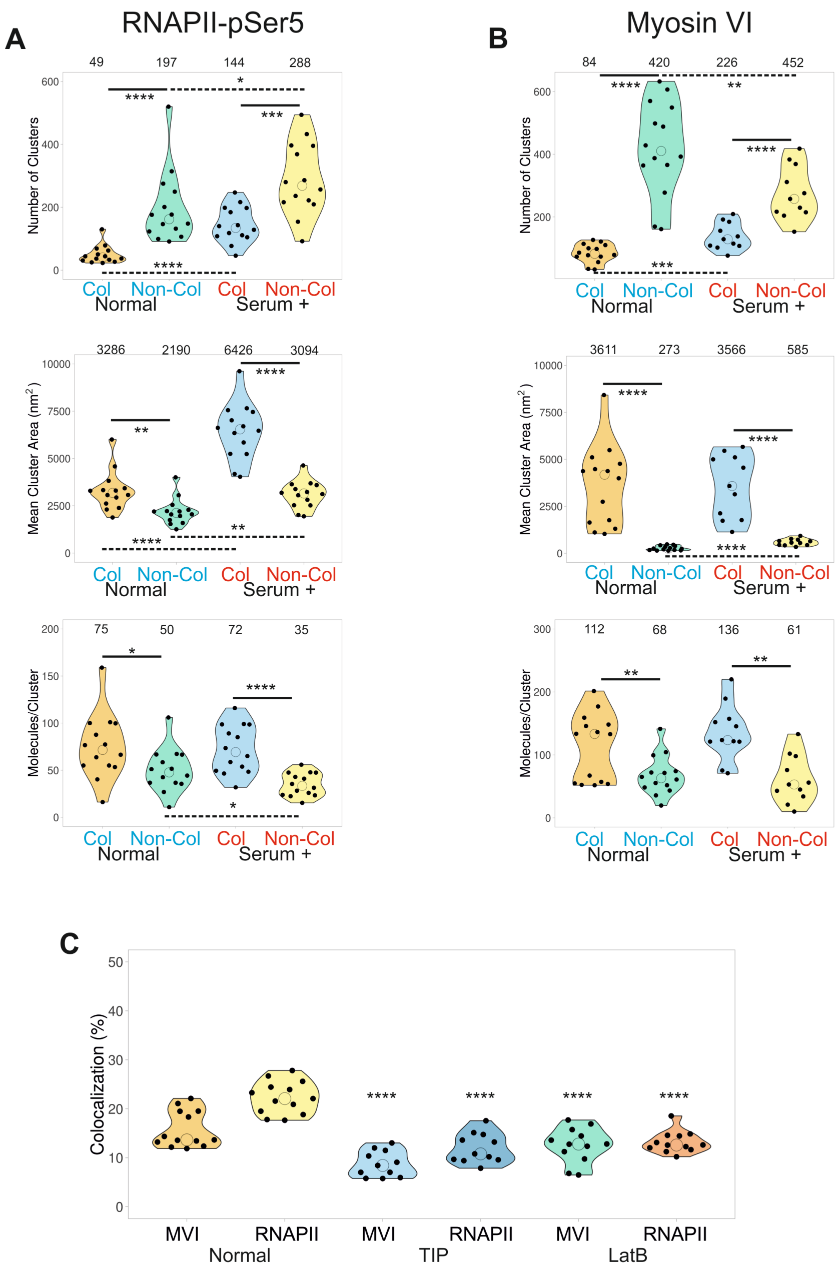


**Supplementary Figure 1 Colocalization analysis of RNAPII with MVI.** (A) Cluster analysis of RNAPII-pSer5 nuclear organisation under normal and serum-treated conditions. The data are broken down into myosin VI colocalized and non-colocalized clusters. Individual data points correspond to the average value for a cell ROI (n=14). The values represent the mean from the ROIs for each condition (Only statistically significant changes are highlighted *p <0.05, **p <0.01, ***p <0.001, ****p <0.0001 by two-tailed t-test compared to normal conditions). (B) Cluster analysis of myosin VI nuclear organisation under normal and serum-treated conditions. The data are broken down in to RNAPII-pSer5 colocalized and non-colocalized clusters. Individual data points correspond to the average value for a cell ROI (n=14). The values represent the mean from the ROIs for each condition (Only statistically significant changes are highlighted **p <0.01, ***p <0.001, ****p <0.0001 by two-tailed t-test compared to normal conditions). (C) Colocalization analysis of myosin VI (MVI) and RNAPII-pSer5 clusters under normal (n=13), TIP- (n=11) and LatB-treated (n=11) conditions. Individual data points represent the percentage of each protein which is colocalized and correspond to the average value for a cell ROI. The values represent the mean from the ROIs for each condition (Only statistically significant changes are highlighted ****p <0.0001 by two-tailed t-test compared to normal conditions for each protein).

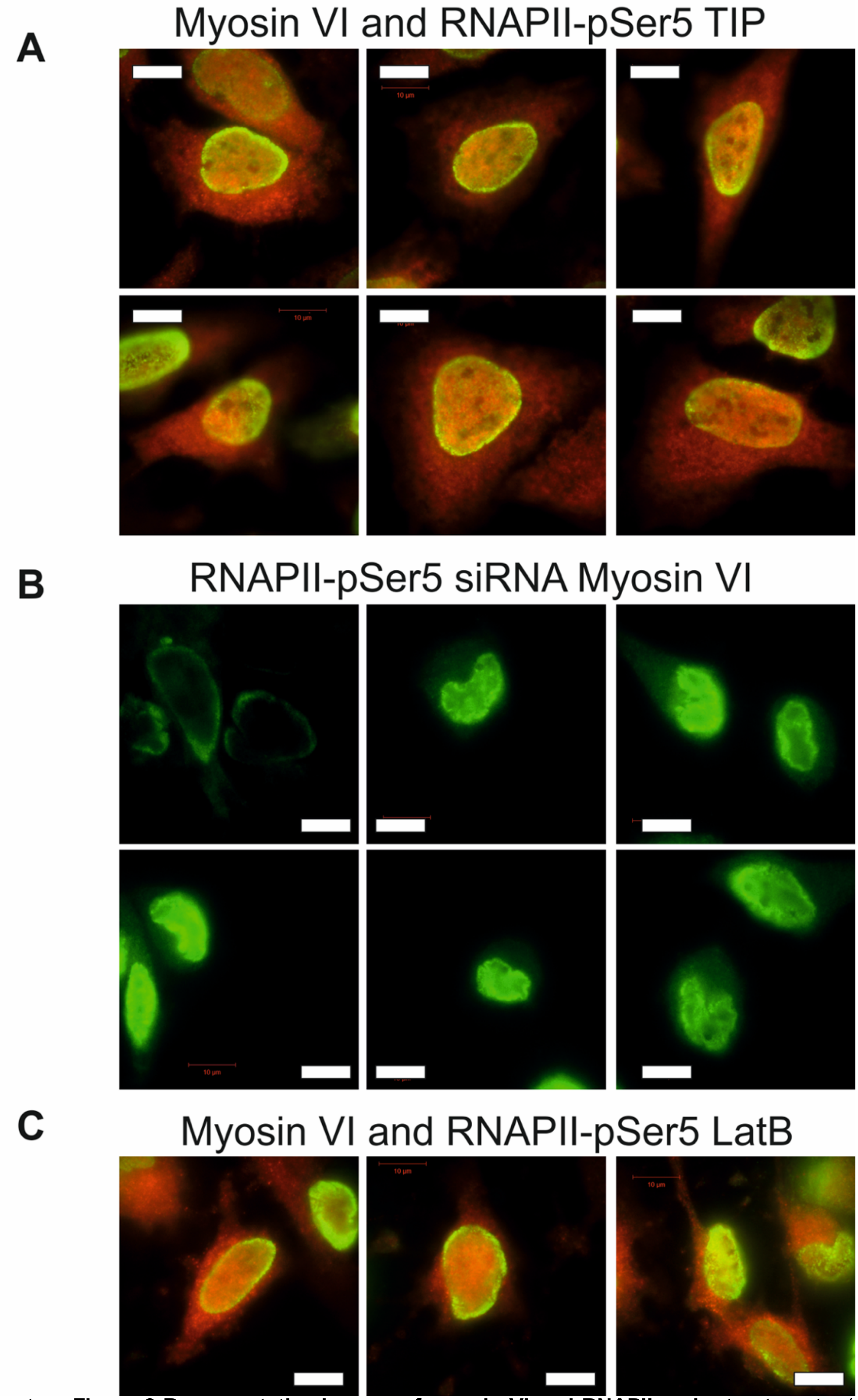


**Supplementary Figure 2 Representative images of myosin VI and RNAPII under treatments.** (A) Widefield Immunofluorescence staining against myosin VI (red) and RNAPII-pSer5 (green) in HeLa cells when treated with TIP. Images were acquired at the mid-point of the nucleus (scale bar 10 μm). (B) Widefield Immunofluorescence staining against RNAPII-pSer5 (green) in HeLa cells following siRNA knockdown of myosin VI. Please see Supplementary Figure 3 for myosin VI controls. Images were acquired at the mid-point of the nucleus (scale bar 10 μm). (C) Widefield Immunofluorescence staining against myosin VI (red) and RNAPII-pSer5 (green) in HeLa cells when treated with LatB. Images were acquired at the mid-point of the nucleus (scale bar 10 μm).

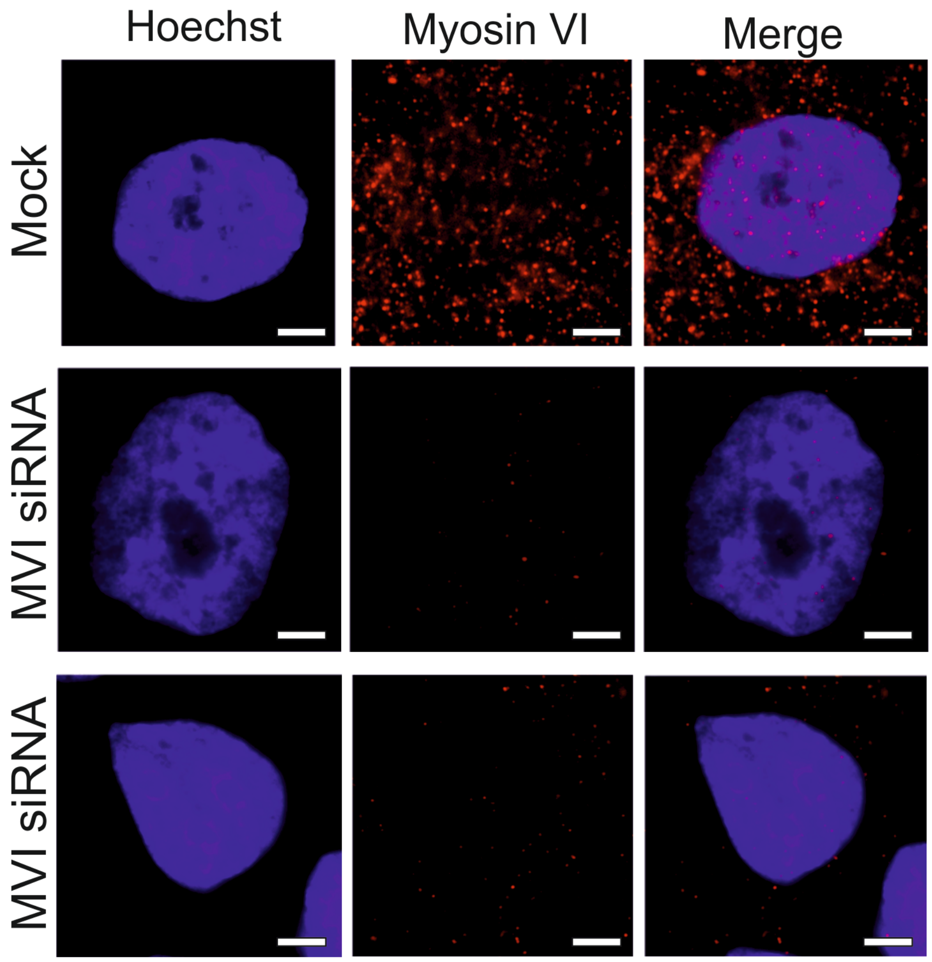


**Supplementary Figure 3 Representative images of myosin VI following siRNA knockdown.** Widefield Immunofluorescence staining against myosin VI (MVI) (magenta) and DNA (cyan) in HeLa cells under mock (control siRNA treatment) and siRNA conditions following 48 hr transfection. Images were acquired at the mid-point of the nucleus (scale bar 5 μm).

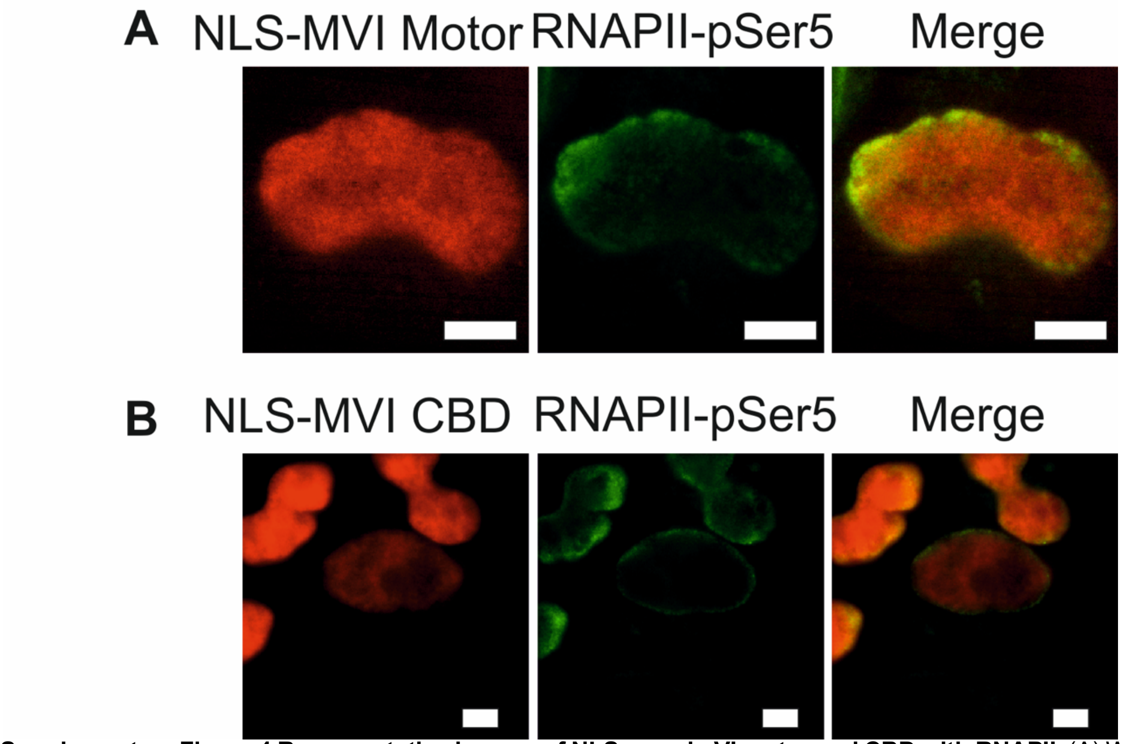


**Supplementary Figure 4 Representative images of NLS myosin VI motor and CBD with RNAPII.** (A) Widefield Halo-TMR and Immunofluorescence staining against NLS myosin VI motor (red) and RNAPII-pSer5 (green) in HeLa cells. Images were acquired at the mid-point of the nucleus (scale bar 5 μm). (B) Widefield Halo-TMR and Immunofluorescence staining against NLS myosin VI CBD (red) and RNAPII-pSer5 (green) in HeLa cells. Images were acquired at the mid-point of the nucleus (scale bar 5 μm).

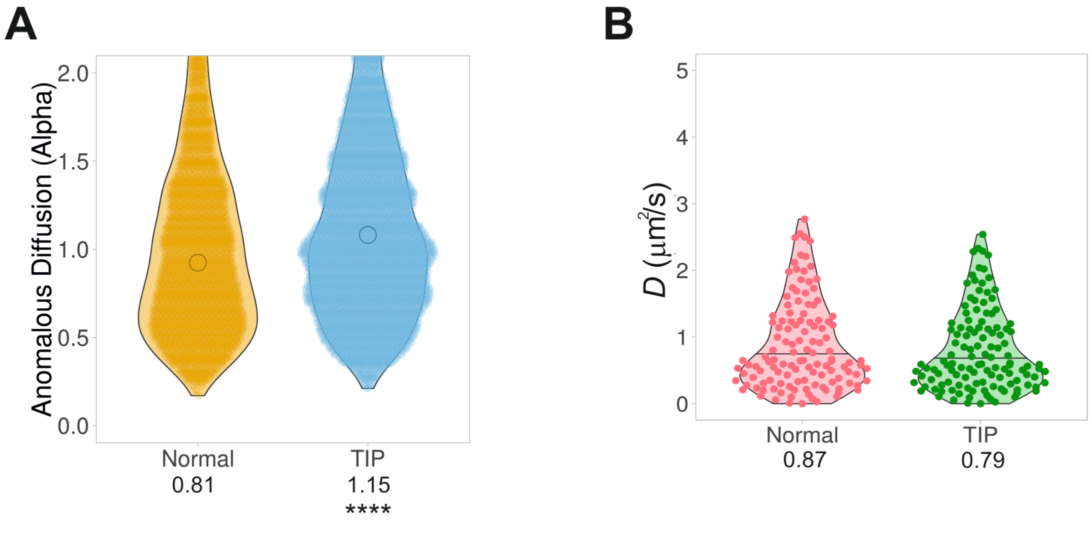


**Supplementary Figure 5 Controls for Live cell single molecule tracking.** (A) Plot of Halo-RNAPII anomalous diffusion alpha value under normal and TIP treated conditions. Values were derived from fitting trajectories to an anomalous diffusion model, as described in methods. Data points correspond to the individual tracks across all cells (n = 100). The values represent the mean from the ROIs for each condition. (****p <0.0001 by two-tailed t-test compared to normal conditions) (B) Plot of SNAP-tag diffusion constants under the stated conditions derived from fitting trajectories to an anomalous diffusion model, as described in methods. Data points correspond to the average value for a cell ROI (n=100). The values represent the mean from the ROIs for each condition. There is not a statistically significant change between the datasets.

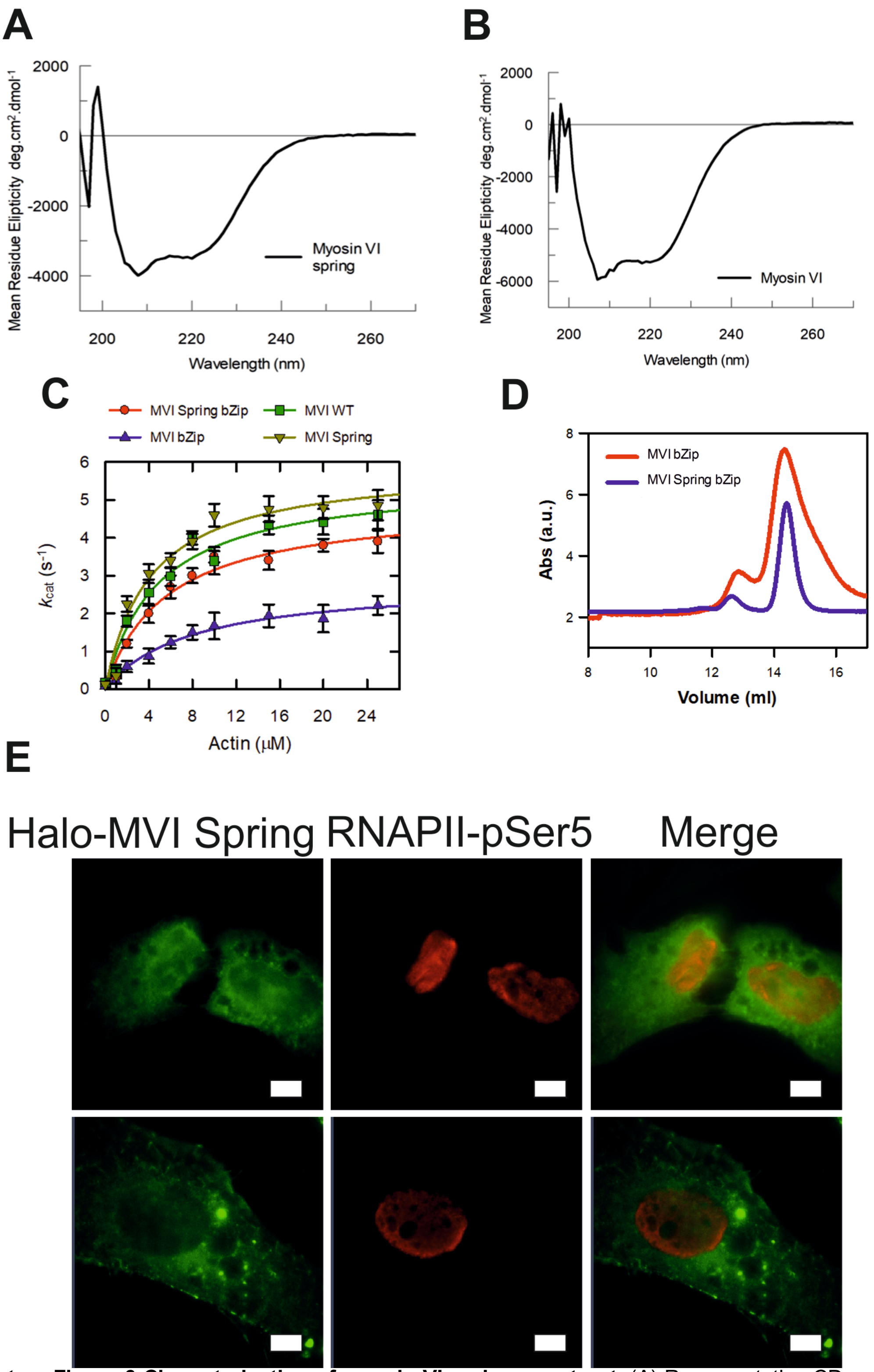


**Supplementary Figure 6 Characterisation of myosin VI spring construct.** (A) Representative CD spectra for recombinant full-length myosin VI spring. (B) Representative CD spectra for recombinant full-length wild type myosin VI. (C) Michaelis-Menten plot of displaying steady-state actin-activated ATPase activity for the myosin VI (MVI) constructs. Error bars represent SEM from three-independent experiments. (D) Representative size exclusion chromatography trace for 1 mg/ml myosin VI bZip and myosin VI spring bZip constructs. (E) Representative widefield Halo-TMR and Immunofluorescence staining against Myosin VI Spring (green) and RNAPII-pSer5 (red) in HeLa cells. Images were acquired at the mid-point of the nucleus (scale bar 5 μm).

**Supplementary Table 1. GO Gene Expression analysis**

| p value (FDR corrected) | Genes in List | Total Genes | Functional Category | GO Term |
|---|---|---|---|---|
| 2.88E-09 | 308 | 3903 | Regulation of cell communication | GO:0010646 |
| 2.88E-09 | 310 | 3952 | Regulation of signaling | GO:0023051 |
| 7.76E-09 | 265 | 3287 | Cell surface receptor signaling pathway | GO:0007166 |
| 7.76E-09 | 280 | 3529 | Regulation of signal transduction | GO:0009966 |
| 7.20E-07 | 244 | 3113 | Intracellular signal transduction | GO:0035556 |
| 1.71E-06 | 269 | 3547 | Response to organic substance | GO:0010033 |
| 2.78E-06 | 169 | 2027 | Regulation of intracellular signal transduction | GO:1902531 |
| 2.78E-06 | 204 | 2561 | Response to external stimulus | GO:0009605 |
| 2.78E-06 | 218 | 2785 | Anatomical structure morphogenesis | GO:0009653 |
| 2.78E-06 | 226 | 2905 | Regulation of localization | GO:0032879 |
| 2.78E-06 | 151 | 1756 | Regulation of cell proliferation | GO:0042127 |
| 2.78E-06 | 344 | 4820 | Regulation of response to stimulus | GO:0048583 |
| 2.78E-06 | 256 | 3382 | Regulation of multicellular organismal process | GO:0051239 |
| 2.78E-06 | 119 | 1286 | Negative regulation of multicellular organismal process | GO:0051241 |
| 2.78E-06 | 64 | 551 | Circulatory system process | GO:0003013 |
| 3.08E-06 | 63 | 542 | Blood circulation | GO:0008015 |
| 3.85E-06 | 161 | 1921 | Locomotion | GO:0040011 |
| 3.90E-06 | 197 | 2474 | Nervous system development | GO:0007399 |
| 4.67E-06 | 99 | 1028 | Regulation of cellular component movement | GO:0051270 |
| 9.19E-06 | 175 | 2165 | Cell proliferation | GO:0008283 |

**Supplementary Table 2. Recombinant DNA.**

| Construct | Source |
|---|---|
| Xenopus pFastBac1 Calmodulin (1-end) | Sellers Lab |
| Human pFastbacHTB MVI (1-1253) | Fili et al 2017 |
| Human pFastbacHTB MVI bzip | Sellers Lab |
| Human pFastbacHTB MVI Spring | Synthetic Gene - This Study |
| Human pFastbacHTB MVI Spring bzip | Synthetic Gene - This Study |
| pHalo-Rpb1 | Darzacq lab |
| pSNAP-C1 | Addgene 58186 |
| pSNAP-Rpb1 | This Study |
| YFP-NLS-R62D Actin | De Lanerolle Lab |
| pLV-Tet0-Halo-NLS-Motor (1-814) | Große-Berkenbusch et al 2020 |
| pLV-Tet0-Halo-NLS-CBD (1060-1253) | Große-Berkenbusch et al 2020 |
| pcDNA3.1 MVI | Synthetic Gene - This Study |
| pcDNA3.1 MVI Spring | Synthetic Gene - This Study |

**Supplementary Table 3. Primers for qPCR.**

| Sequence | Use |
|---|---|
| CTGCTGCAGGAACCTCTCAT | VASP For |
| CTTCCTGGGGAGAATGTGG | VASP Rev |
| GACAACTCCTGGTTGGGAGA | FOXJ2 Pro. For |
| GGAGGTCCTACTTGGGGAAG | FOXJ2 Pro. Rev |
| AGGGGGAATCTCTAGGCAAA | PTEN Pro. For |
| TGCATTCGCTCTTTCCTTTT | PTEN Pro. Rev |
| GAATGGTTTGTGGCTCAGGT | PTEN Int. For |
| CCCACTGGGTTGAAATATGG | PTEN Int. Rev |
| GGTGCCTGAGAAGAGGTGAG | EGR3 ex2 For |
| CCATGTGGATGAATGAGGTG | EGR3 ex2 Rev |
| TCACGTACCACACACACACG | EGR3 Int. For |
| TCCGGGTCTGAAACTACCTG | EGR3 Int. Rev |
| CCCCTGCTTCTTCTCAGTTG | IER3 Pro. For |
| AAAAGATGCACGGATTGGAG | IER3 Pro. Rev |
| CTCACCTTCCTCCTCCTCCT | TNNT1 Int. For |
| GTCCAGCAGAGACTGGAACC | TNNT1 Int. Rev |
| GGCTCACCCTTGCTGATGCT | Myc For |
| GCTCTGGGCACACACATTGG | Myc Rev |
| GCTTTCCTGCCTCACCATTA | FOXJ2 Int. For |
| AAGGCCAAGTCAGATGAAGC | FOXJ2 Int. Rev |
| GGAGAAGAACAGCACAACCT | VASP RT-qPCR For |
| CCCTCTGTAGGTCCGAGTAAT | VASP RT-qPCR Rev |
| TGGTCAAGGCAGAACAGAAG | TNNT1 RT-qPCR For |
| CCCATGTAGTCAATGTCCAGAG | TNNT1 RT-qPCR Rev |
| CTCGGATGGAGGTTACTCTTTC | INHA RT-qPCR For |
| CACCAGCCATGGGATTAAGA | INHA RT-qPCR Rev |
| TCAGCCCAGTGTTGTCATTAG | CDC42BPA RT-qPCR For |
| TCTTTCCGACTTGCATGGATAA | CDC42BPA RT-qPCR Rev |
| GGCTCCTTTGAAGTGGAGTAATA | EGR3 RT-qPCR For |
| CACAGGAGAAGTAACGCTAACA | EGR3 RT-qPCR Rev |
